# Phasic and tonic pain serve distinct functions during adaptive behaviour

**DOI:** 10.1101/2025.02.10.637253

**Authors:** Shuangyi Tong, Timothy Denison, Danielle Hewitt, Sang Wan Lee, Ben Seymour

## Abstract

Pain drives self-protective behaviour, and evolutionary theories suggest it acts over different timescales to serve distinct functions. Whilst phasic pain provides a teaching signal to drive avoidance of new injury, tonic pain is argued to support recuperative behaviour, for instance by reducing motivational vigour. We test this hypothesis in an immersive virtual reality EEG foraging task where subjects harvested fruit in a forest: some fruit elicited brief phasic pain to the grasping hand, and this reduced choice probability. Simultaneously, tonic pressure pain to the contralateral upper arm was associated with reduced action velocities. This could be explained by a free-operant computational framework that formalises and quantifies the function of tonic and phasic pain in terms of motivational vigour and decision value. Additional analyses examined the relationship between model parameters, subjective ratings, and physiological and neural responses. Overall, the results show how tonic and phasic pain subserve distinct objective motivational functions that support harm minimisation during ongoing adaptive behaviour.

## INTRODUCTION

Despite its unpleasantness, pain serves a useful function. This is most clearly the case for phasic pain, where newly incoming nociceptive signals warn of impending tissue damage, elicit rapid defensive responses, and drive learning that reduces future chances of encountering harm. In this context, function can be objectively measured through conditioned responses and avoidance behaviour, which quantify pain in terms of its motivational value (Fields, 2006; Seymour, 2019). However, the adaptive function of tonic pain has been much harder to quantify. In ecological and ethological contexts, tonic pain is generally considered to serve protective and recuperative functions. For instance, in the context of injury, it has been proposed that tonic pain can aid recovery by reducing motivational vigour and hence suppress unnecessary activity until healing occurs (Bolles and Fanselow, 1980; Wall, 1979; Walters and Williams, 2019; Seymour et al., 2023). The goal of this study was to test whether we can formally and quantifiably dissociate these two distinct motivational functions of pain.

One of the challenges in studying adaptive functions of pain is the difficulty of embedding experiments within ecologically meaningful contexts. To solve this, we designed an immersive foraging task using virtual reality (VR), in which humans search a forest to collect fruits from the low-lying bushes at varying heights. A foraging paradigm provides a robust, free-operant framework that captures the core components of adaptive behaviour: it is goal-directed, involves complex movement, and requires the learning of an optimal strategy to maximise rewards. This allows us to computationally dissociate how different types of pain influence the control of action.

Our first hypothesis was that phasic pain provides a distinct valuation signal that updates the value of specific actions within complex environments. In our task, this was implemented by associating specific fruit (distinguishable by colour) with a brief electrical stimulus to the grasping hand, emulating thorns. In our computational model, this was defined as an aversive utility term incorporated into the state-action value evaluation process. We predicted that this computational mechanism would manifest behaviourally as a reduction in choice probability for pain-associated targets and an increase in ‘choice distance bias’ (the willingness to travel further for pain-free options). Neurally and physiologically, we predicted that these aversive values would be tracked by skin conductance responses (SCRs) and the amplitude of nociceptive event-related potentials (ERPs), specifically the N1-P2 complex (Favero et al., 2023).

Second, we hypothesised that tonic pain acts as a coefficient modulating the trade-off between opportunity cost and vigour cost, thereby serving a recuperative function. To test this in Experiment 2, we delivered continuous tonic pressure to the non-dominant arm via an inflated cuff to emulate a background state of injury. Within our free-operant framework, tonic pain was modelled as a weighting factor that shifts the optimal balance toward reduced energy expenditure. Because the stimulus was applied to the non-task limb, we specifically predicted a global reduction in motivational vigour—operationalised as decreased movement velocities and foraging rates—rather than a direct mechanical impairment. By applying this formal computational approach, we move beyond exploratory observations to provide a rigorous, mechanism-based explanation for how distinct pain states adaptively govern choice and action.

## RESULTS

### Experiment 1: Phasic pain avoidance in a free-operant foraging task

Twenty-five subjects performed a free-operant instrumental foraging task to accrue reward points and avoid pain. The task was embedded into a fully immersive VR context, in which subjects physically moved in a flat open space. A virtual boundary of 4 m × 4 m was displayed when the subject approached the edge of the area. Within the VR context, this space resembled a flat forest (Fig. 1), with trees and vegetation locations randomly generated. The task was structured into one-minute blocks. At the beginning of each block, virtual fruits of varying heights and locations were generated within the virtual space. The spatial coordinates of subjects’ heads and hands were tracked, requiring them to physically move towards the fruit, pick it up by reaching with their dominant hand, clicking, and holding the button on the handset. Only the dominant hand was enabled in the virtual environment. Subsequently, subjects could drop the fruit into one of the baskets to earn points. Subjects were told that the total points accrued would be rewarded with an incentive of up to £10. All fruits used in both experiments were pineapples, as their spiky shape naturally matches the pricking pain induced by the Wasp pain stimulation electrodes.

**Figure 1.**
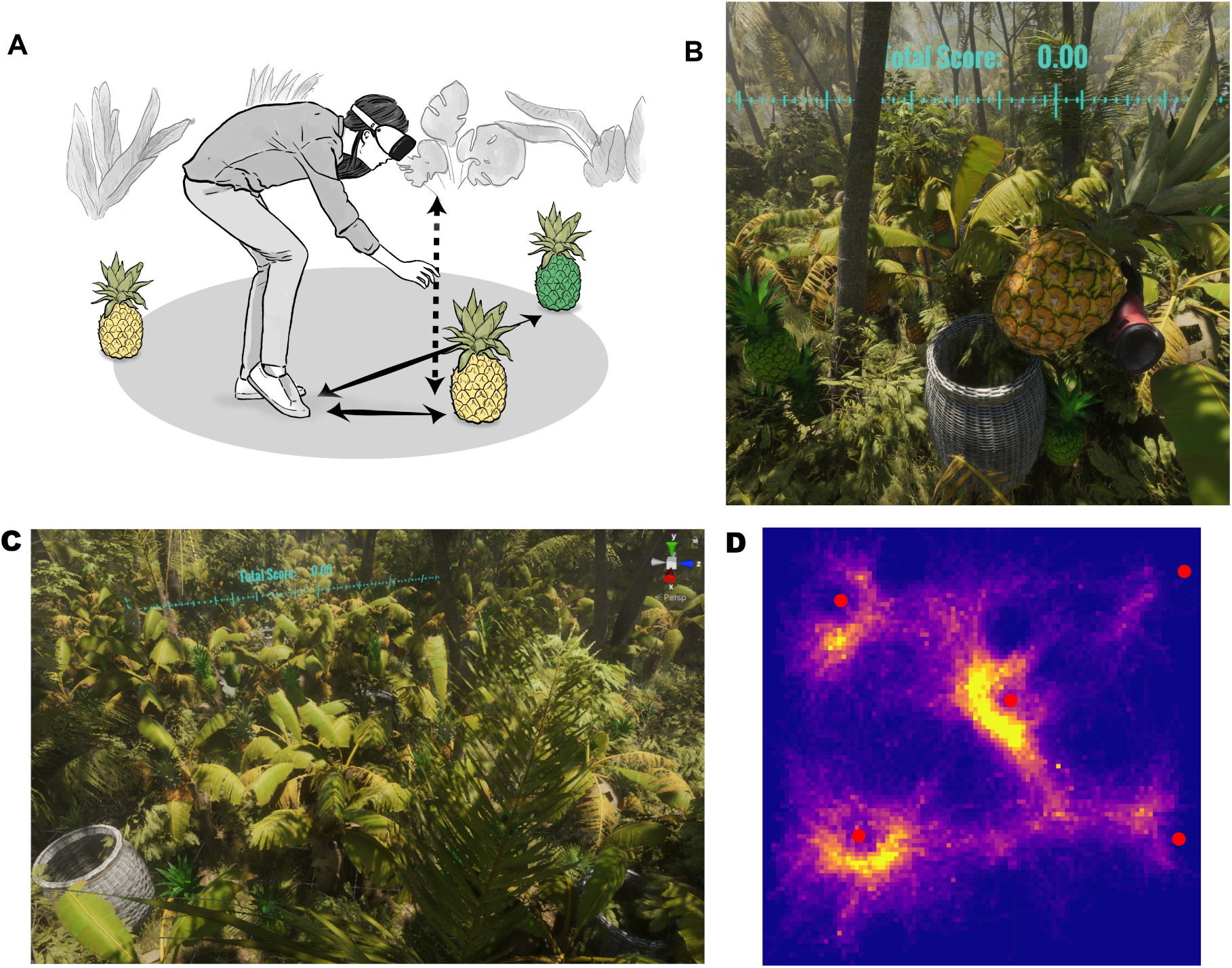
Task description. (**A**) An illustrative figure depicting the foraging task. (**B**) Subjects’ view (left eye) in the task. (**C**) A perspective view of the task environment. (**D**) An aggregated top-view heat map of head trajectories of all subjects’ data. Red dots denote baskets’ preset locations.

Pineapples were categorised into two visually distinguishable types: 50% were green and 50% were yellow. The green pineapple was aversive, and picking one up immediately elicited a brief painful cutaneous electric stimulation to the proximal medial side of the middle finger of the (grasping) dominant hand. In contrast, picking up a yellow pineapple was always pain-free. The intensity of the electric stimulation was held constant within each block. Experiment 1 consisted of 20 blocks, with the initial 10 blocks designated for practice purposes with a fixed order of stimulation intensities. The subsequent 10 blocks used randomised pain stimulation intensities across five different levels, determined by a pain calibration procedure. After completing the 10 training blocks, all subjects were aware that picking up green pineapples resulted in painful shocks, as confirmed through verbal questioning. However, subjects did not know the exact pain level of a new block until they tried to pick up one green pineapple. Only data from the latter 10 non-training blocks were analysed.

#### Avoidance increases with increasing phasic pain intensity

We found that the probability of choosing a fruit decreased as the pain associated with it increased. For this analysis, both aversive choice probabilities and subjective pain ratings were estimated at the block level. Figure 3A shows an approximately linear decrease in aversive choice probability as a function of visual analogue scale (VAS) rating of pain stimuli, whereby the aversive choice probability is defined as the number of painful fruits collected divided by the total number of fruits collected.

#### Additional cost of effort associated with movement

Next, since movement itself may carry a small cost related to time and effort (incurring opportunity costs), we investigated whether there is a trade-off relationship between moving distances and pain intensity (i.e., is it worth moving further to avoid a certain amount of pain). We measured the distance from the fruit to the subject at the first fixation on that fruit within the search window, as estimated from eye-tracking data. The window began at the most recent fruit collection with a valid fixation-to-pickup measurement, or at block onset if none was available; skipped collections did not advance its start. We found a significant correlation between the egocentric distance differences between painful and non-painful fruits, and the VAS ratings. As shown in Figure 3B, the vertical axis represents the ‘choice distance bias’, calculated as the difference between the average egocentric distance to non-painful fruits and the average egocentric distance to painful fruits within each block. The egocentric distance is the fruit distance relative to the participant. This metric was computed to test whether subjects would trade off physical effort for pain avoidance; specifically, a positive bias indicates that subjects were willing to bypass closer painful fruits to reach more distant pain-free ones. As hypothesised, we found that as the pain intensity (VAS) of the aversive fruits increased, this distance bias grew significantly, confirming that subjects exerted greater movement effort to avoid higher levels of pain.

#### Choice-based computational modelling in RL framework

As noted above, these initial regression results were analysed at the block level. Within each one-minute block, each pickup, which could result in an immediate shock, can be considered an individual trial. By analysing the data at the trial level, we can extract more detailed information and provide a parameterised behavioural description for each individual. This approach laid the groundwork for a robust, objective measure of phasic pain value within an ecologically valid context. In each trial, despite the presence of numerous lower-level controls required to complete the action, the primary decision involved choosing which fruit to pick up. We simplified the problem by assuming that there exists a set of decision points *S* as states, where the subject took an action at a state *s* ∊ *S*—choosing to pick up a fruit—and deterministically arrived at another state *s* ′ ∊ *S* if the block had not concluded. Based on the initial regression results, we assumed that the reward was a linear combination of an internal reward *r* for each fruit, a negative utility function *u* representing the phasic pain value, and an effort cost function *d*. Therefore, the Bellman optimality equation for the action-value function was

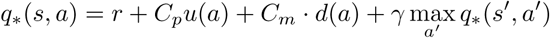

where *C_p_* and *C_m_* were the parameters to be fitted for each subject.

As fruits were randomly distributed in our task, previous foraging studies suggested that animals may adopt a suboptimal greedy policy, opting for the closest fruit rather than solving the complex travelling salesman problem for a marginally better optimal solution (Anderson, 1983; Jeon et al., 2023). To initially demonstrate the basic effects, we further eschewed learning and future planning by setting the discount factor *γ* = 0. Figure 2 shows the conceptual diagram of the computational framework setup. One of the key simplifications was assuming the existence of a set of decision points in this free-operant context. To fit the model, we again utilised the eye-tracking data, setting the decision point as the first time point when the subject saw a new fruit. Nevertheless, the task’s available actions were partially observable, and subjects could explore for more actions if they were not satisfied with the currently available ones. This exploration action was not modelled in our formulation. To proceed with model fitting, we retrospectively identified the fruits that were picked up and selected only those time points when the fruits chosen afterwards were first seen as the decision points. We also kept track of the previously seen fruits, storing them in a memory queue, so the decision involved choosing the best option from these items in memory. Figure 3(C, D) shows the parameters fitted to the subjects’ behavioural data. This shows the negative utility associated with pain and vertical and horizontal components of movement, and it illustrates how the computational modelling framework can be used to quantify behaviour in a free-operant VR context.

**Figure 2.**
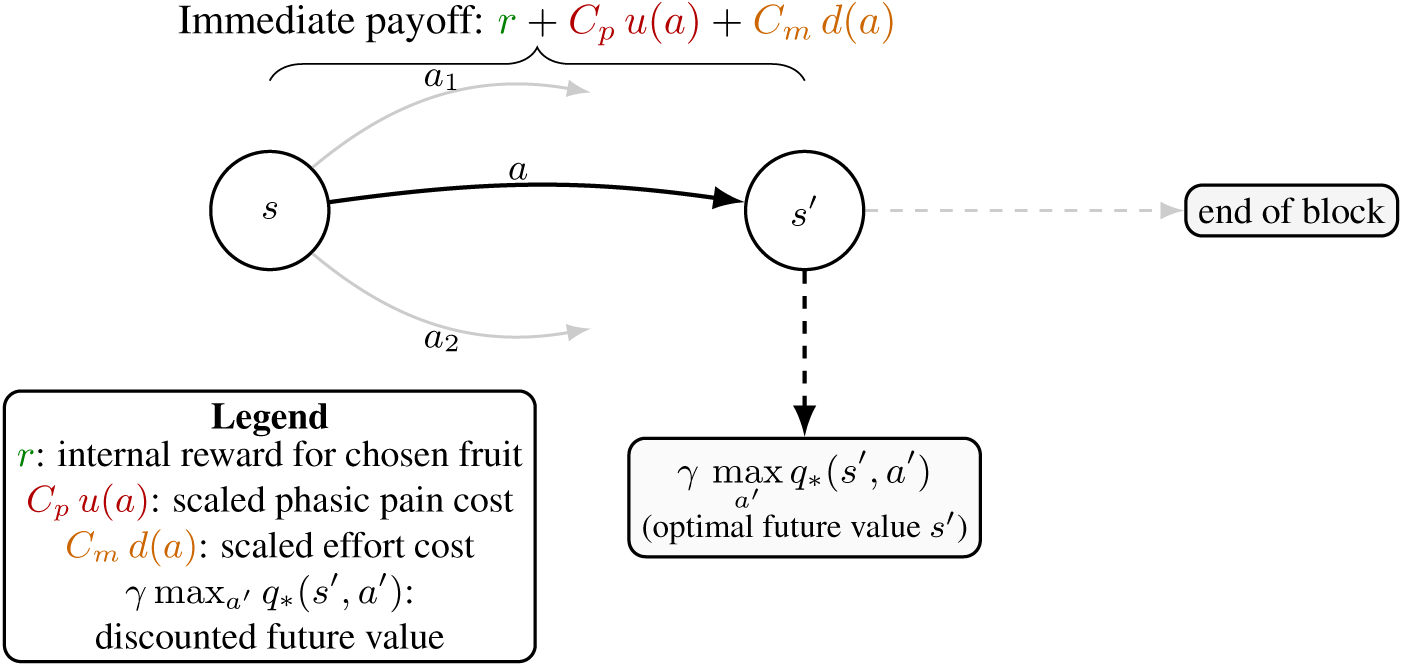
Illustration of the computational setup for a single trial within a block. From state *s*, the subject chooses an action *a* (picking a fruit), deterministically transitions to *s^′^* if the block continues, receives an immediate payoff decomposed into reward, pain, and effort, and then accrues discounted future value.

#### Skin conductance responses and the interpretation of pain decision values

The model provides an estimate of pain’s influence on choice, raising the question of how this decision value relates to the pain participants report. Decision values were correlated with subjective pain ratings (Fig. 3G), but this relationship leaves open whether they capture the same aspects of pain or whether the choice-derived measure provides additional information about its aversive value. We therefore sought physiological evidence that could help clarify this distinction. Skin conductance responses (SCRs) offered a way to investigate the question at different stages of the task: fixation on pain-associated fruit can accompany evaluation of the anticipated cost of selecting it, whereas the shock delivered at pick-up is directly linked to receiving pain. These contexts motivated us to compare the associations of decision values and ratings with fixation- and shock-evoked SCRs, seeking clues about the relationship between prospective valuation and subsequent pain experience.

**Figure 3.**
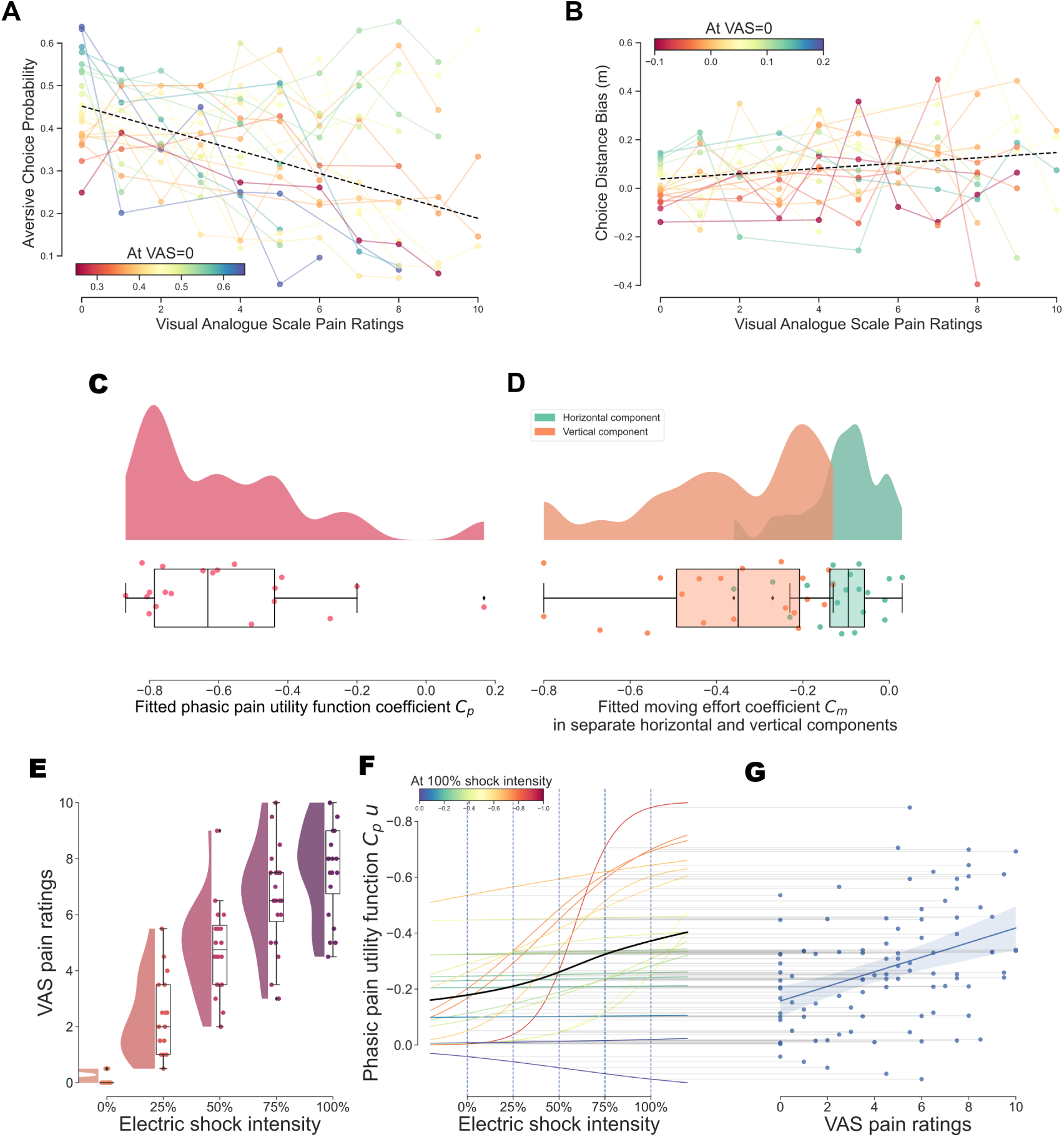
Experiment 1 behaviour and modelling results. (**A**) Each coloured line represents one single subject data with the colour reflecting the value in pain-free condition (VAS=0). The dashed black line’s slope and intercepts are fixed effect estimates from a linear mixed model. The slope’s estimate shows an inverse relationship between pain choice probability and VAS rating, *β* =-0.026, 95%*CI*[-0.039, −0.014], *t*(21.2) =-4.43, *p <* .001. (**B**) As described in (A), the dashed line’s slope is the fixed effect’s estimate of choice distance bias, and it shows a positive relationship between the choice distance bias and VAS ratings, *β* = 0.011, 95%*CI*[0.001, 0.020], *t*(53.05) = 2.29, *p* = .026. (**C**) The negative phasic pain coefficients (*M* = −0.583*, SD* = 0.254) showed the model captured the aversiveness of phasic pain stimuli in this free-operant decision-making task, *t*(19) = −9.994*, p <* .001. (**D**) The moving effort coefficient *C_m_* comprised a horizontal component *h* (*M* = −0.110*, SD* = 0.093) and a vertical component *v* (*M* = −0.376*, SD* = 0.205). A two-sided paired *t*-test of the within-subject difference *h v* showed a lower fitted effort cost to move horizontally than vertically, *t*(19) = 6.37*, p <* .001. (**E**) VAS ratings at different electric shock intensities. (**F**) Phasic pain utility values as a function of electric shock intensity. Each coloured line represents one subject’s fitted curve. Black line is the average over all subjects. (**G**) A mixed model accounting for hierarchical data (random intercepts and slopes per subject) showed significant correlation between VAS pain ratings and model estimated phasic pain values, *β* = *−*0.024, 95%*CI*[*−*0.039*, −*0.009], *t*(17.38) = −3.41, p = .003.

Figure 4A shows SCRs aligned to fixation and pick-up events. To estimate the magnitude of potentially overlapping responses, we fitted the skin conductance time series using event regressors convolved with a canonical response function (Bach et al., 2010). The resulting block-level coefficients for fixation and pick-up of pain-associated fruit were analysed in separate, unweighted multilevel linear mixed-effects models, without selecting coefficients according to their first-level statistical significance. Shock-evoked SCR coefficients were positively associated with subjective pain ratings (Fig. 4C). The associations between fixation-evoked SCR coefficients and either ratings or decision values, and between shock-evoked SCR coefficients and decision values, remained inconclusive in this primary analysis (Fig. 4B,C).

**Figure 4.**
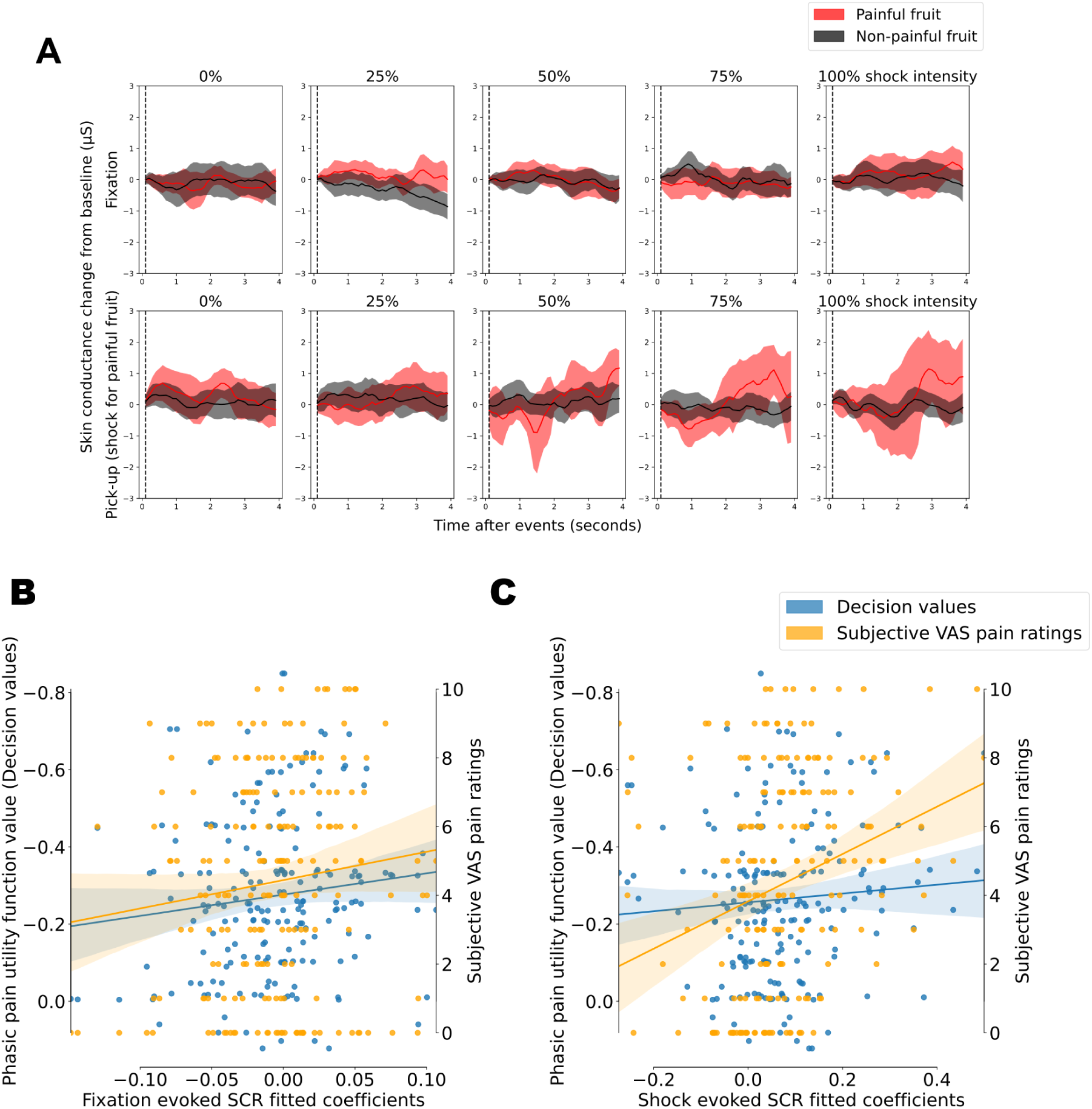
Skin conductance responses and their associations with subjective pain ratings and model-derived decision values in Experiment 1. (**A**) SCRs aligned to fixation and fruit pick-up events for painful and non-painful fruit. Shaded areas indicate 95% confidence intervals. (**B**) In the primary unweighted mixed-effects analyses, the associations of fixation-evoked SCR coefficients with decision values (*β* = −0.008, 95% CI [-0.019, 0.003], *t*(6.79) = −1.72, *p* = .130) and subjective ratings (*β* = 0.006, 95% CI [-0.006, 0.019], *t*(16.20) = 1.13, *p* = .274) were inconclusive. (**C**) Shock-evoked SCR coefficients were positively associated with subjective ratings (*β* = 0.039, 95% CI [0.015, 0.063], *t*(17.02) = 3.38, *p* = .004); the association with decision values was inconclusive (*β* =-0.010, 95% CI [-0.042, 0.022], *t*(8.08) = −0.73, *p* = .487). The models included 195 fixation and 196 shock coefficients from 20 participants. Slopes express the change in the fitted SCR coefficient per one standard deviation of the predictor. Block coefficients were not selected by first-level statistical significance; the remaining exclusion criteria are described in Methods. Regression lines in panels B and C describe the pooled data; the reported statistics are from the mixed-effects models.

The shock–rating association was the only association supported across both experiments by all three approaches: the unweighted analysis and supplementary inverse-variance mixed-effects analyses using `metafor` and `brms`, which incorporated the estimated uncertainty of the block-level coefficients (Supplementary materials, Skin conductance sensitivity analyses). Shock-evoked coefficients were also associated with decision values in Experiment 2, while evidence for fixation–rating associations depended on the analysis method. The present analyses do not resolve whether decision values and ratings reflect the same or partly distinct aspects of pain; further study is needed to determine whether their physiological associations can help distinguish these interpretations.

### Experiment 2: Modulation of free-operant foraging by tonic pain stimulation

In the second experiment, we introduced a tonic pain condition by applying an inflatable blood pressure cuff around the non-dominant arm (Graven-Nielsen et al., 2017). We reduced the number of phasic pain levels from five in the first experiment to three (no pain, low pain, and high pain). These three phasic pain levels were then combined with two fixed tonic pain conditions (with and without tonic pain) to form a factorial design experiment with a total of six different combinations. Similar to experiment 1, the experiment was divided into one-minute blocks, and the pain conditions were fixed within each block. The six combinations were initially presented in a fixed order as training blocks, then the order was randomised and repeated three times, resulting in a total of 24 blocks. To obtain a neural measure of pain behaviour, we introduced concurrent EEG recording alongside the VR set-up.

#### No effect found for tonic pain on phasic pain

We found no significant modulation of phasic pain ratings by tonic pain (Fig. 5(A, B)). We found a basic decrease in aversive choice probability, in terms of the probability of selecting a painful fruit, as in experiment 1. The average aversive choice probabilities were similar in conditions with and without tonic pain (Fig. 5(C-E)), providing evidence that punishment sensitivity was not affected by the presence of tonic pain.

**Figure 5.**
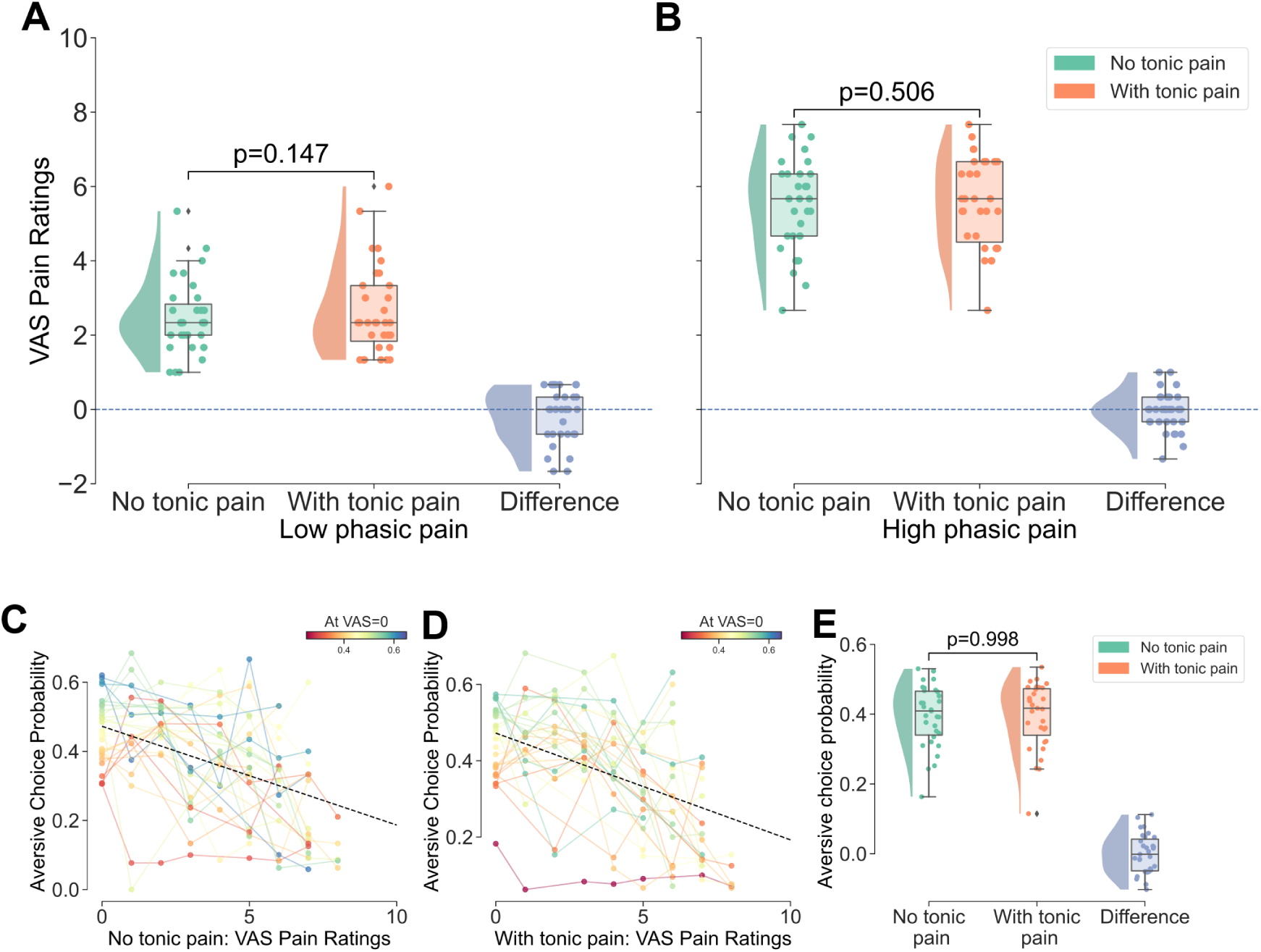
Effects of tonic pain on phasic pain ratings and aversive choice probabilities. (**A, B**), Two-way repeated measures ANOVA showed no statistically significant difference in subject’s ratings of phasic pain intensity as a result of tonic pain in low (A) or high (B) phasic pain conditions, F(1,30)=2.01, p=.167, nor was there a significant interaction effect between tonic pain and phasic pain, F(1, 30)=0.87, p=.357. (**C, D**), Similar linear mixed model fitting results for aversive choice probability. No tonic pain condition: *β* = *−*0.029, 95%*CI* = [*−*0.038*, −*0.020]*, p <* .001, intercept = 0.473. Tonic pain condition: *β* =*−*0.028, 95%*CI* = [*−*0.035*, −*0.021]*, p <* .001, intercept = 0.472. (**E**), Aversive choice probability averaged over phasic pain conditions. Two-way repeated measures ANOVA showed significant effect for phasic pain (including no pain condition), *F* (2, 60) = 36.87*, p <* .001, but no effect for tonic pain, *F* (1, 30) = 0.00*, p* = .998. The interaction effect between tonic and phasic pain was also not significant *F* (2, 60) = 0.07*, p* = .930.

In the neural data, we found that phasic pain intensity modulated the amplitude of phasic pain ERPs, as would be expected. However, we found no significant tonic-pain effect on ERP amplitude or tonic-by-phasic intensity interaction (Fig. 6), consistent with the behavioural results. We focused our neural analysis of phasic pain on ERPs as phasic stimuli are well characterised by these time-locked evoked potentials. Complementary analyses of total event-related and ERP-subtracted induced power at Cz are reported in the Supplementary materials (section: Event-related time–frequency and induced oscillatory responses to phasic pain stimuli; Figs. S4–S7). High–low phasic pain contrasts were significant in total power under both tonic conditions; in induced power, this contrast reached significance without tonic pain but not with tonic pain. Neither the direct tonic-condition contrasts nor the tonic-by-phasic intensity interaction reached significance in either power representation.

**Figure 6.**
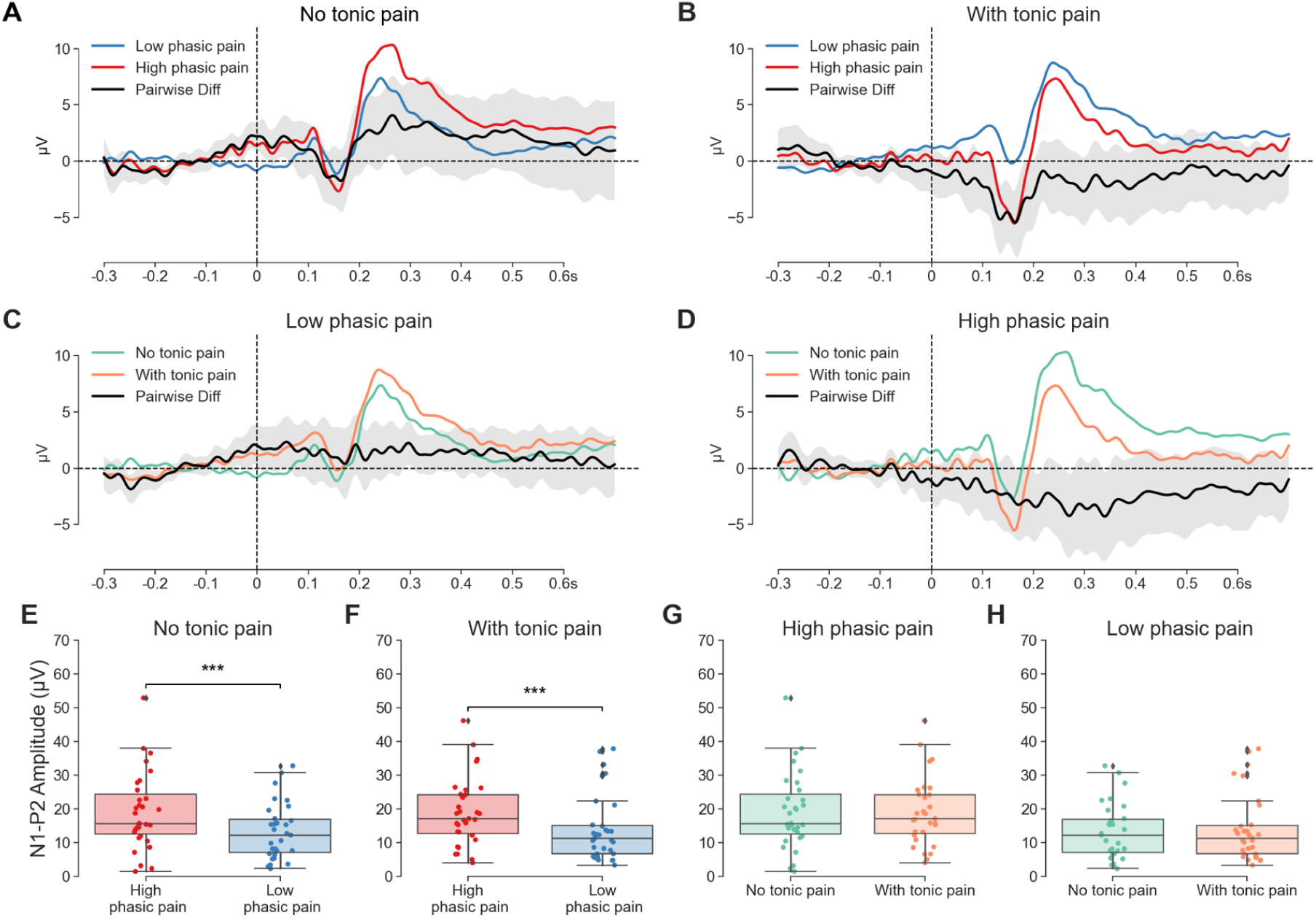
Phasic pain ERPs in different pain conditions. (**A, B**) Phasic pain ERP comparison in the same tonic pain conditions. (**C, D**) Phasic pain ERP comparison in the same phasic pain conditions. (**E, F**) High phasic pain induced a significantly higher N1-P2 amplitude with or without tonic pain. Shaded areas represent the 95% confidence interval across participants. (**G, H**) Tonic pain stimulation does not show a significant effect in phasic pain ERP’s N1-P2 amplitude. Two-way repeated measures ANOVA showed the effect for phasic pain was significant, *F* (1, 30) = 35.42*, p <* .001. The effect for tonic pain was not significant, *F* (1, 30) = 0.30*, p* = .589, and the interaction effect was also not significant, *F* (1, 30) = 0.58*, p* = .454.

#### Tonic pain reduced motivational vigour

By analysing the motion tracking data, we found that tonic pain was associated with a reduction in task-related movement velocities (Fig. 7(A, C)) and an associated reduction in fruit collection rates (Fig. 7B).

**Figure 7.**
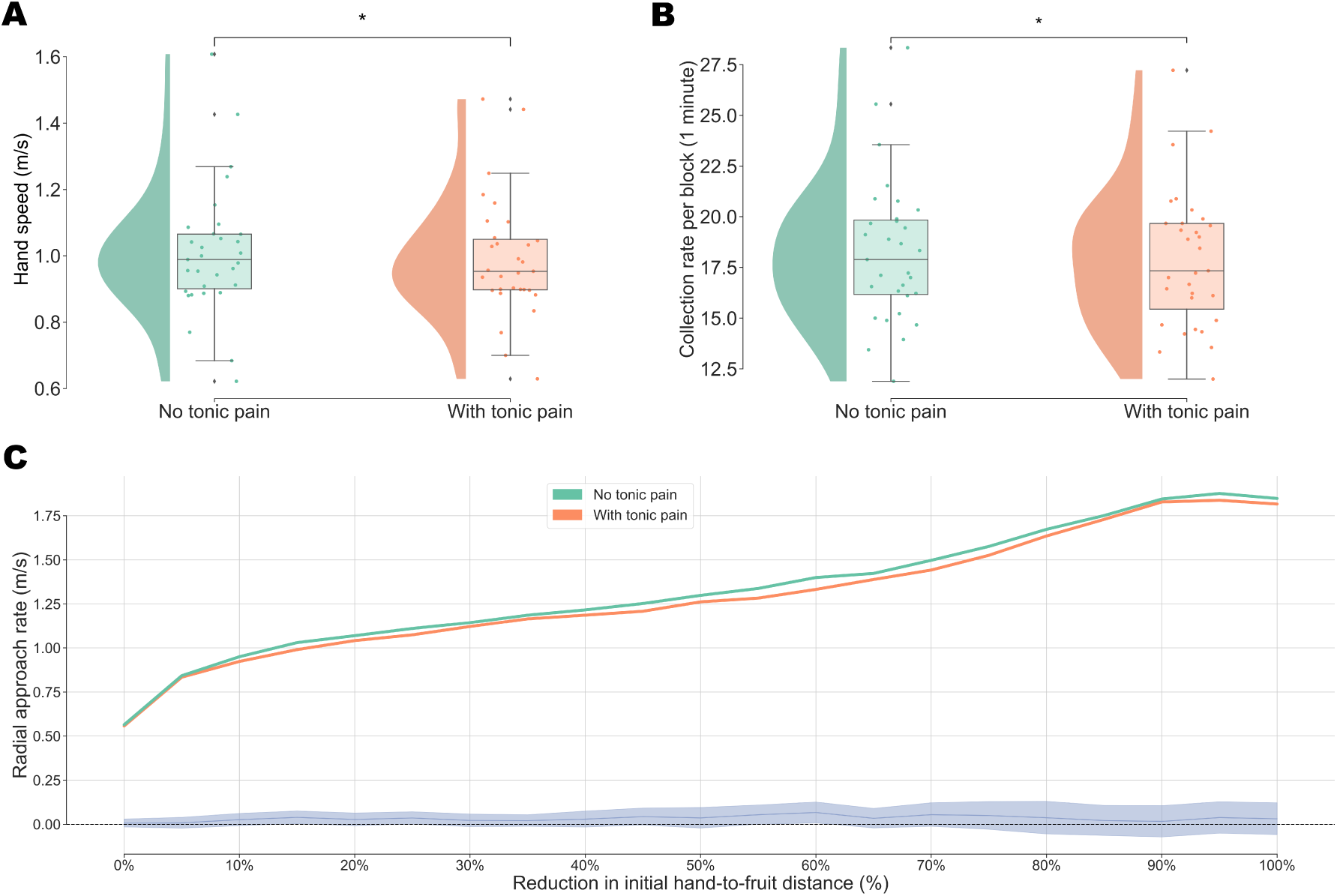
Tonic pain reduced action velocity. (**A**) Average hand speed. A one-tailed paired t-test was conducted to evaluate whether the hand speed was faster without tonic pain (*M* = 1.01*, SD* = 0.19) than with tonic pain (*M* = 0.99*, SD* = 0.18), *t*(30) = 2.09, *p* = .023, with a small to moderate effect size *d* = 0.37. (**B**) The average fruit collection rate over the one-minute block. A one-tailed paired t-test was conducted to evaluate whether the collection rate was higher without tonic pain (*M* = 18.22*, SD* = 3.45) than with tonic pain (*M* = 17.91*, SD* = 3.35), *t*(30) = 1.93, *p* = .031, with a small to moderate effect size *d* = 0.35. (**C**) Estimated radial approach rate of the hand towards the fruit as a function of progress in reducing the initial straight-line hand-to-fruit distance. Rates were calculated from interpolated crossing times, with pickup marking completion. Shaded area is the 95% confidence interval of the pairwise difference mean.

To quantify this effect in terms of motivational vigour, we extended the computational model to accommodate vigour effects. The model presented for experiment 1 does not consider time, and fitting the model to the collected data requires acausal simulation (see Methods, Model description and fitting section below). For experiment 2, we took account of the temporal factor and fitted the model causally with high temporal resolution. Niv et al. (2007) proposed that the delay between consecutive actions performed by animals can be formulated as a trade-off between opportunity costs and vigour costs. This framework was developed based on earlier works in average-reward reinforcement learning, showing that this trade-off approach maximises average reward (Niv et al., 2005). In this formulation, animals choose an action pair (*a, τ*). The first, *a*, is the common action in discrete decision-making models, and the separate *τ* represents the time delay between two actions. We built upon this model to suit the requirements of our study. Similar to the model in experiment 1, our action *a* represents the choice of which fruit to pick up. To account for the different effort costs incurred for each fruit, we again used a distance-based effort cost term *C_m_ · d*(*a*), as in experiment 1. Additionally, a time delay between actions is required to model vigour. We assumed that subjects estimated the time delay by estimating their speed *V_speed_* and calculated the delay based on the distance to the fruit and the estimated speed. Hence, for a decision time point state *s* ∊ *S*, if the effort cost is inversely proportional to the time delay, the optimal differential value of an (*a, τ*) pair is

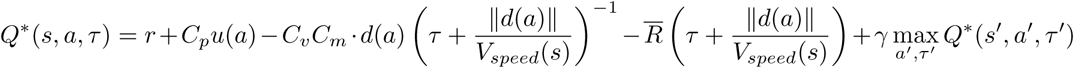

where *τ* is the additional delay accounting for the time waited before committing to execute the action. *s^′^ ∊ S* is the deterministic subsequent state after choosing (*a, τ*). *Q^∗^* is the optimal differential state-action value function. *C_v_* is the vigour constant that scales the vigour cost term. *R̅* is the average reward and assumed to be constant (Fig. 8).

**Figure 8.**
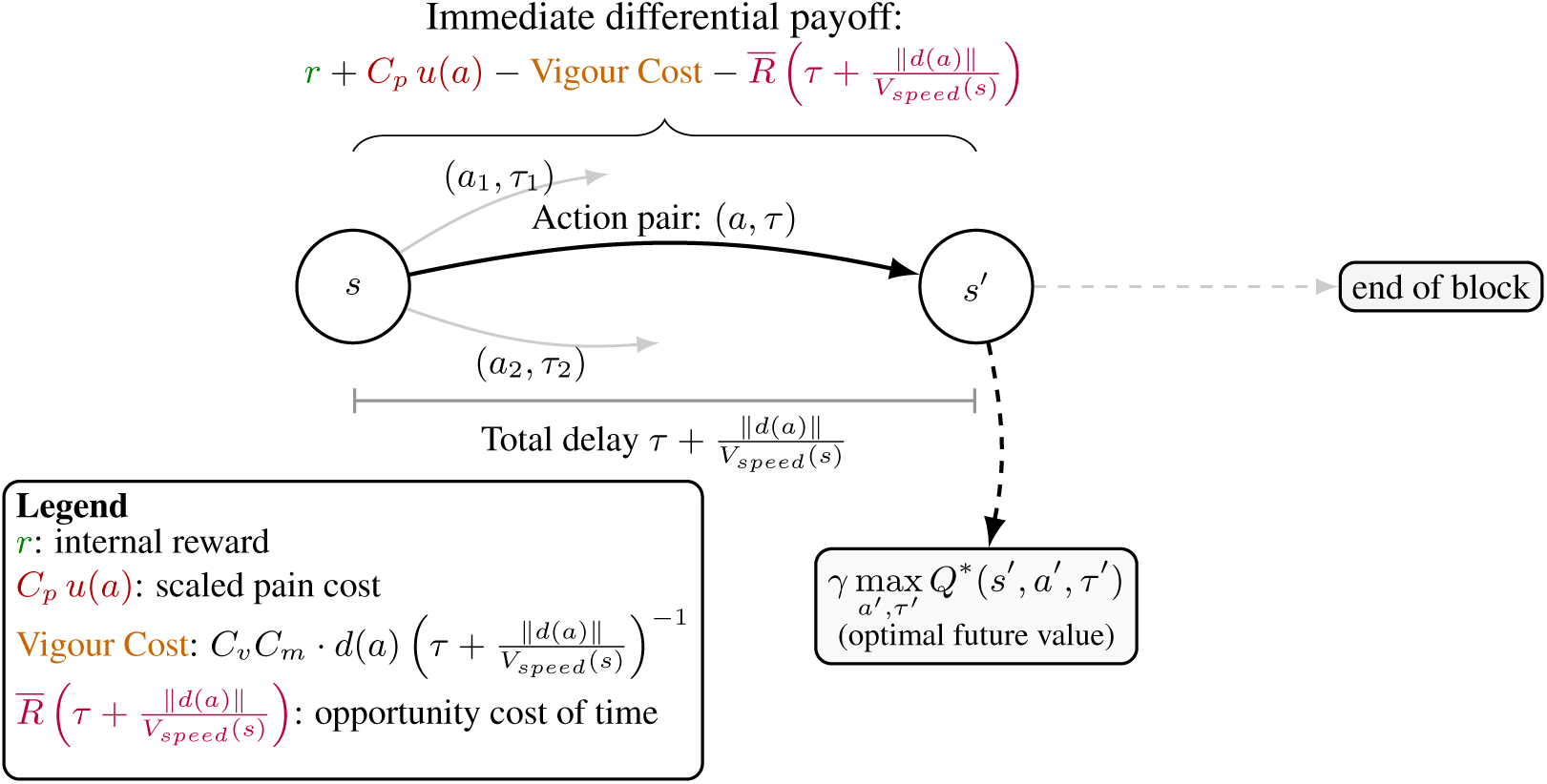
Computational setup for Experiment 2 incorporating vigour and opportunity costs. The subject chooses a fruit *a* and a latency *τ*. The total delay 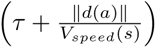 penalises the payoff via both a vigour-dependent effort cost and the opportunity cost of time.

We fitted vigour constants *C_v_* and other parameters for no tonic pain and with tonic pain conditions separately. We found the fitted vigour constants were significantly higher in tonic pain conditions (Fig. 9A). We further fitted the model with additional separate vigour constants for different levels of phasic pain. Consistent with the fitting results over tonic pain conditions only, tonic pain showed a strong effect on fitted vigour constants, while phasic pain did not show a significant effect (Fig. 9B). However, it is worth noting that optimal vigour constants were nonunique and the condition contrast was sensitive to the selection among equally fitting parameter combinations. The *C_v_* fitting result presented here is the first minimum encountered in the ascending joint-grid search. We explain the rationale for using such method in Methods, Model description and fitting. Limited identifiability may partly reflect the structure of the available choice sets.

**Figure 9.**
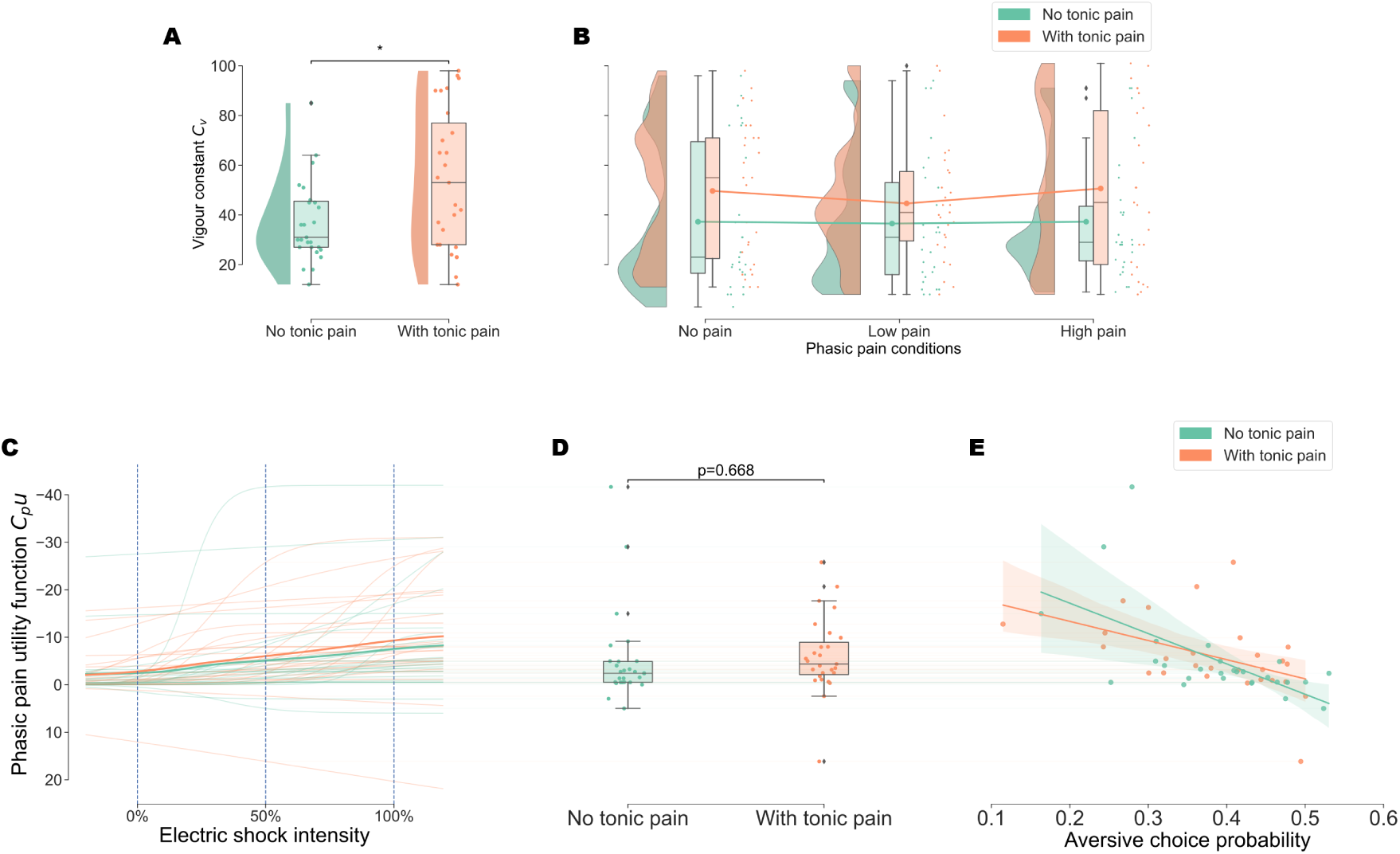
Experiment 2 model-fitting results. (**A**) Fitted vigour constants were greater in tonic pain conditions (*M* = 54.04*, SD* = 27.41) than no tonic pain conditions (*M* = 38.44*, SD* = 18.15), *t*(26) = 2.28, *p* = .031. (**B**) Separate vigour constants fitting for each phasic pain conditions. Repeated measures ANOVA showed the effect for tonic pain was significant, *F* (1, 26) = 4.91*, p* = .036, but the effect for phasic pain was not significant, *F* (2, 52) = 0.18*, p* = .832. The interaction effect was not significant, *F* (2, 52) = 0.20*, p* = .818. (**C**) Phasic pain utility function curves. Solid lines are the average values in each condition. (**D**) Phasic pain utility function values at 50% electric shock intensity. A paired t-test was conducted and found no significant differences between no tonic pain (*M* = *−*5.10*, SD* = 9.48) and with tonic pain (*M* = *−*6.02*, SD* = 7.90) conditions, *t*(26) = *−*0.43, *p* = .668. (**E**) Phasic pain utility function values at 50% electric shock intensity plotted against aversive choice probability. Both conditions show a significant correlation between the fitted model values and behavioural choice probability. In blocks without tonic pain, *R*^2^ = 0.356, *F* (1, 25) = 13.80, *p* = .001; in tonic pain conditions, *R*^2^ = 0.221, *F* (1, 25) = 7.07, *p* = .013.

#### A unified model for tonic and phasic pain for decision-making in free-operant task

This new decision model therefore includes both phasic pain and vigour costs, by which the tonic pain effects can be expressed as change in vigour. Experiment 2 replicated the phasic pain punishment sensitivity effect shown in experiment 1. Using this model, the fitted phasic pain parameters also showed no significant differences in punishment sensitivity (Fig. 9(C-E)), in line with the behavioural results presented above. The vigour constants and phasic pain function fitting results illustrate how we can combine two dissociable effects of pain motivation in a single model, consistent with the results from empirical regression.

#### Characterisation of neural activity during tonic pain

To probe the neural activity associated with the modulation of motivational vigour by tonic pain, we performed LMM-based time-frequency analysis on the EEG data. The data were epoched based on key decision points. Linear mixed models (LMM) were employed to account for random effects across subjects and to control for motion artefacts by including instantaneous head movement speed as an additional predictor. We found that alpha band power over parietal and centroparietal regions (Pz, P4, P8 and CP6), as well as beta power at P8, was negatively associated with the presence of tonic pain at the uncorrected level (Fig. 10A). Although these electrode-wise associations did not survive Bonferroni correction, the alpha cluster survived cluster-based permutation correction (*p*_cluster_ = .006); the beta association did not (*p*_cluster_ = .082).

**Figure 10.**
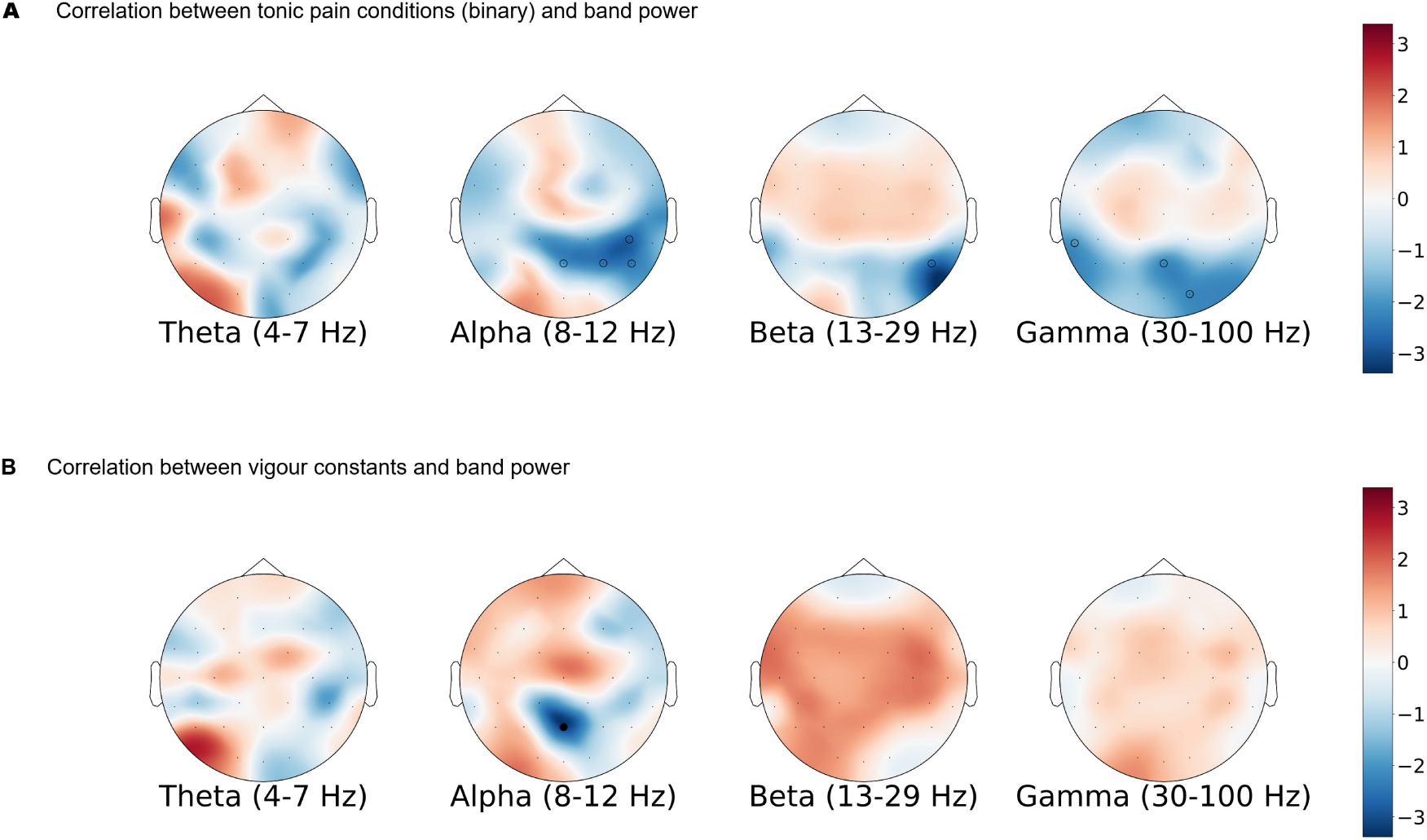
(**A**) Topography of t-values from the LMM, with binary tonic pain condition as the independent variable and EEG band power 0–0.5 s after the decision point as the dependent variable. In both panels, empty circles indicate uncorrected *p <* .05, and solid circles indicate *p <* .05 after Bonferroni correction across the 32 electrodes within each frequency band. Tonic pain was associated with lower alpha power at Pz (*t* = *−*2.07), P4 (*t* = *−*2.50), P8 (*t* = *−*2.42) and CP6 (*t* = *−*2.74), and lower beta power at P8 (*t* = *−*2.56). Lower gamma power was also observed at TP9 (*t* = *−*1.96), Pz (*t* = *−*2.14) and O2 (*t* = *−*2.20). None of these electrode-wise associations survived Bonferroni correction. The alpha cluster comprising Pz, P4, P8 and CP6 survived cluster-based permutation correction (*p*_cluster_ = .006), whereas the beta P8 cluster (*p*_cluster_ = .082) and gamma clusters (TP9: *p*_cluster_ = .291; Pz and O2: *p*_cluster_ = .089) did not. (**B**) Topography of t-values from the LMM, with continuous fitted vigour constants as the independent variable and EEG band power 0–0.5 s after the decision point as the dependent variable. A negative association with alpha band power was found at midline parietal electrode Pz (*t* = *−*3.21, uncorrected *p* = .001), which survived Bonferroni correction (*p*_Bonferroni_ = .042) but did not reach the cluster-corrected significance threshold (*p*_cluster_ = .059).

To investigate how tonic pain impacts vigour, the vigour constants separately fitted to tonic pain conditions (Fig. 9A) were used as predictors of EEG band power. We found that the alpha band power over the midline parietal region (Pz) was negatively correlated with the vigour constants and survived Bonferroni correction (*p*_Bonferroni_ = .042), but fell just short of the cluster-corrected significance threshold (*p*_cluster_ = .059; Fig. 10B).

## DISCUSSION

The experiments show that tonic and phasic pain serve different motivational functions during adaptive behaviour, in line with ecological and evolutionary theories of pain (Bolles and Fanselow, 1980; Walters and Williams, 2019). Specifically, our findings point towards phasic pain providing a punishment teaching signal that directs avoidance through value-based learning, balancing the cost of future harm alongside potential reward. This is supported by the observation that increasing phasic pain intensity significantly reduced choice probability and increased distance bias between choices, whereby participants were willing to travel further to reach a pain-free fruit. In contrast, we found that tonic pain reduces motivational vigour, which supports energy conservation and recuperation in the context of bodily damage. This claim is directly evidenced by the reduction in task-related movement velocities and fruit collection rates during tonic pain blocks. The experiments are the first to show that these two functions can be formally distinguished and quantified during ongoing behaviour. By utilising a free-operant RL computational framework, we were able to dissociate these roles: phasic pain was quantified as a generally negative utility term affecting choice values, while tonic pain was formalised as a change in vigour constants that were significantly higher (increasing delays between actions) in tonic pain condition, under the joint-grid selection rule described in Methods. This illustrates how pain simultaneously acts in different ways to serve self-protection.

One notable aspect of our results is that we did not see interactions between tonic and phasic pain at either the behavioural or neural level. Behaviourally, we observed that average aversive choice probabilities remained similar regardless of the presence of tonic pain, with no significant interaction effect on punishment sensitivity. Furthermore, our model-fitting confirmed that tonic pain did not significantly modulate the fitted phasic pain utility values. There are two contexts in which these might be predicted. First, in ‘conditioned pain modulation’ paradigms (Kennedy et al., 2016), a tonic pain stimulus is sometimes seen to reduce both the perceived intensity and the cortical evoked responses to phasic pain stimuli delivered somewhere else on the body (Höffken et al., 2017; Enax-Krumova et al., 2020). Although we utilised concentric ‘Wasp’ electrodes designed to selectively activate nociceptive A-delta fibres (Inui et al., 2002), and confirmed that the resulting ERPs (N1-P2) were significantly modulated by phasic intensity, we observed no such attenuation by tonic pain. Indeed, neither subjective pain ratings nor the N1-P2 amplitude showed a significant modulation by the tonic pressure pain stimulus. In contrast, our results were more compatible with a trend in the other direction.

This finding raises the (testable) question as to what extent conditioned pain modulation depends on the task setting: notably, here we utilise an active in contrast to a passive task, which is known to be an important distinction in other contexts of coping behaviour (Bandler et al., 2000). In an active behavioural context, the brain may prioritise the fidelity of phasic pain as a teaching signal for learning, potentially overriding modulation seen in passive CPM paradigms. A second context in which phasic-tonic interactions are possible is in punishment sensitivity, where it is proposed that tonic pain signals increase vulnerability to new phasic pain insults (Seymour et al., 2023). Punishment sensitivity is seen in terms of anxiety-like responses to novel threats in animals (Crook et al., 2014; Lister et al., 2020), and also in decision-making tasks in people with chronic pain (Mancini et al., 2024). However, we do not see an increase in the punishment value of phasic pain in relation to tonic pain here.

It is worth noting that, while we found no clear behavioural or ERP evidence that tonic pain modulated phasic pain responses, this does not establish that phasic pain processing is independent of tonic pain. Supplementary analyses identified a significant high–low phasic pain contrast in induced power without tonic pain but not with tonic pain; however, direct interaction tests were nonsignificant for both total event-related and induced power. These findings warrant caution because the free-operant task allowed participants to avoid high-pain outcomes, leaving limited and unequal high-pain trial counts. The subtracted ERP estimate may therefore retain non-phase-locked activity and noise, limiting the separation of evoked and induced activity.

These two contexts highlight the potential difference between ratings and choice as measures of aversiveness, a concept that is sometimes referred to as experienced utility and decision utility (Kahneman and Tversky, 1984), or disliking and unwanting (the negative version of the liking-wanting distinction (Dayan and Seymour, 2009; Berridge et al., 2009), or put colloquially, ‘what you say’ versus ‘what you do’). In our task, we explored this possibility by comparing the associations of decision values and pain ratings with fixation-evoked and phasic pain-evoked (shock) SCRs; the association between shock-evoked SCRs and ratings was consistently supported, but the findings did not resolve whether the two measures have distinct physiological correlates. Further research is needed to determine whether these measures capture distinct aspects of the overall aversiveness of pain, highlighting the value of combining subjective ratings with model-based measures of its motivational impact on behaviour to address this question.

The experiments show how probing objective motivational functions of pain is facilitated by VR. Conventionally, pain motivation is studied using desktop tasks in which decisions and physiological responses are made by a seated subject, responding via a keyboard with pain stimuli applied to a distant site. VR provides several useful advances. First, the ability to apply pain stimuli in a way that is appropriate to the action, i.e. phasic pain stimuli to a grasping hand, providing sensorimotor congruency. Second, the ability to embed pain within a free-operant and free-moving physical task allows evaluation of full-body responses to a topographic stimulus, and it is this in particular that allows assessment of motivational vigour. Third, the provision of a multisensory immersive environment entirely transforms a pain task into a naturalistic setting that matches the experience of pain in real-life. The importance of naturalistic contexts is well documented in fear learning, although less well studied for pain (Mobbs et al., 2015).

However, the use of VR requires an analysis framework that accommodates the complexity of free-operant and free-moving nature of behaviour. This challenge can be overcome with a computational approach and judicious use of simplifying assumptions where appropriate. The free operant model exploits a theoretical framework developed for animal studies that derives action delays from a trade-off between vigour cost and opportunity cost (Niv et al., 2007). This model has been applied to human studies (Nair et al., 2023), albeit in a much simpler setting involving perfect information and a single instantaneous action. In our task, the environment was partially observable and evolved as subjects interacted with it. By utilising eye-tracking to extract an abstract observation space and a modified version of the model that enabled simulation at high temporal resolution, we demonstrated that it is possible to predict free-operant behaviour in real-time. We use motivational vigour here for the regulation of effort and pace in goal-directed behaviour, informed by accounts of competing motivations in pain (Fields, 2006). Movement speed and collection rate describe its behavioural expression, while the fitted *C_v_* provides a choice-based operationalisation of vigour-related movement costs. Its motivational interpretation is a modelling assumption that depends on the validity and adequacy of the model; neither approach uniquely identifies motivation. The fit profiles motivate developing future tasks with richer choice sets to better constrain individual vigour constants (Fig. 11). Richer choice sets may also increase search, memory and planning demands, requiring additional modelling assumptions and parameters.

A concern that is sometimes raised in pain experiments is to what extent the impact of pain on other behavioural and cognitive processes might be interpreted as an attentional phenomenon (e.g. distraction, or cognitive load), and therefore not specific to pain per se (Moont et al., 2010), specifically the reduction in vigour observed here. One argument to this point posits that attention is an intrinsic part of what pain is, and therefore it does not make sense to consider pain as an operational system without attention (Van Damme et al., 2010): i.e. tonic pain may mediate vigour effects by using distraction as a functional mechanism, and therefore attention is a core part of pain, and not a confound. One way to test this would be to add an additional condition comprising a tonic salient but non-painful stimulus (such as a continuous salient auditory stimulus), although this introduces other confounds related to the incommensurability of such dissimilar types of stimulation. Attentional arguments have also been proposed in CPM paradigms, but here it has been noted that CPM (and diffuse noxious inhibitory control) operates at least partly at a spinal/brainstem level, at ‘lower’ levels than those typically associated with attention (Youssef et al., 2016; Kucharczyk et al., 2023).

It is important to acknowledge that the signal-to-noise ratio in both our physiological and neural recordings is lower than that typically observed in conventional, stationary laboratory experiments (Gramann et al., 2011). This is primarily due to the motion artefacts inherent in an immersive and active virtual reality environment. Whilst we utilised robust cleaning and artefact-correction methods (Klug and Gramann, 2021), the elevated noise floor may limit our capacity to detect more subtle neural effects or interactions. These challenges highlight a critical area for future methodological research, particularly in the development of hardware and signal-processing tools designed to isolate neural signals during complex, mobile behavioural tasks.

Despite the greater levels of noise in EEG recordings due to our highly mobile task, an LMM-based approach (that incorporates head-movement speed into the model) suggested reduced alpha and beta power over parietal and centroparietal scalp regions during tonic pain, with cluster-corrected support for the alpha association. The direction and scalp distribution of the alpha pattern are consistent with previous reports of contralateral parietal and somatosensory alpha suppression during tonic pain (Ferracuti et al., 1994; Huishi Zhang et al., 2016; Dowman et al., 2008); the weaker beta association is also compatible with reported beta suppression associated with tonic stimulus intensity (Schulz et al., 2015). When examining how the neural correlates of tonic pain related to motivational vigour, the negative association between alpha power and fitted vigour constants at Pz survived Bonferroni correction but fell just short of the cluster-corrected significance threshold. This tentative association is consistent with the sensory- and task-related components of pain-induced alpha suppression (Hu et al., 2013) and the broader involvement of alpha oscillations in sensory integration and attention (Palva and Palva, 2011), providing a plausible link to processes relevant to motivational vigour without establishing a specific neural mechanism.

It is also important to consider the spatial configuration of the stimuli used in this study. Phasic pain was delivered to the grasping hand to maintain spatial congruency with the virtual fruit, ensuring a coherent nociceptive feedback signal for the interactive task. Additionally, tonic pain was applied to the contralateral arm to prevent mechanical interference with motor execution, which would have occurred if pressure were applied to the ipsilateral limb used for grasping the controller. Whilst this design promotes spatial congruency and avoids mechanical confounds, future studies might explore how these effects generalise across different body parts, for which VR experiments serve as a promising tool to test relevant hypotheses (Hewitt et al., 2026).

Finally, the methodological framework has clinical applicability as a way to objectively measure pain behaviour in chronic pain. For instance, while conventional clinical trials often rely on Quantitative Sensory Testing (QST), which can deviate from real-world scenarios, our immersive VR task provides an ecologically valid and behaviourally sensitive evaluation of pain as a function of diagnosis or treatment. In addition, it could be implemented using real-time analysis to provide a feedback signal for closed-loop interventions, such as interactive VR rehabilitation or deep brain stimulation.

## METHODS

### The free-operant foraging task

Foraging theory is a broad and well-studied subject within animal behaviour research. Various foraging study paradigms have been established in both laboratory and natural environments (Constantino and Daw, 2015; Charnov, 1976; Lihoreau et al., 2011). A natural foraging task, while simple for a healthy human to perform, encapsulates key components relevant to pain research. Firstly, foraging is goal-directed; subjects actively search for food under varying levels of motivation, allowing for the examination of the interaction between motivation and pain. Secondly, a natural foraging task involves full-body movement. Given that many types of pain are closely linked to movement and exercise, motion capture technology—a fundamental component of immersive VR—allows for precise recording and analysis of movement under different pain conditions. Finally, foraging involves exploring the environment and learning an optimal strategy to maximise rewards, naturally linking pain to cognitive functions. Motivated by these considerations, we designed the free-operant foraging task aimed at simulating realistic foraging behaviours.

At the beginning of each one-minute block, a total of 150 virtual pineapples of varying heights from 0.33 to 1 m were randomly generated in a circle centred around the participant with a diameter of 6.67 m. Five identical baskets were placed within the space. Spatial locations of trees and vegetation were generated using the game engine’s default tree painting tool (Unity Technologies, San Francisco, US).

While the colour association (green for painful, yellow for pain-free) was not counter-balanced across subjects, any inherent aversive value of green pineapples (e.g. as ‘unripe’ fruit) is expected to have a minimal confounding effect on the analysed data. In associative learning frameworks, while mild prior biases may influence initial value estimations, extensive training with a highly salient unconditioned stimulus (e.g. phasic pain) rapidly updates these values, driving them toward an asymptote determined entirely by the explicit task contingencies (Rescorla and Wagner, 1972; Sutton and Barto, 2018). Because participants underwent extensive training (10 blocks in Experiment 1 and 6 blocks in Experiment 2) to establish the explicit pain associations prior to the analysed sessions, the observed avoidance behaviour was predominantly driven by the learned phasic pain contingencies rather than baseline colour preferences.

We used the HTC Vive Pro Eye headset and controllers in combination with Valve’s SteamVR platform to track user motion in our Unity-based virtual environment. The system employs Lighthouse tracking, which uses infrared laser-emitting base stations and IR photodetectors to determine the position and orientation of tracked devices. Motion data from the headset and controllers were accessed in Unity via the SteamVR Unity plugin, which maps physical movements to virtual objects in real time. This setup enables accurate and low-latency motion representation in the virtual space, supporting immersive interaction during the experiment (Sansone et al., 2022). Data acquisition in Unity occurred at the rendering frame rate. The average delay between consecutive frames was 22.49 ± 5.82 ms.

Eye-tracking acquisition, quality control, and an exploratory gaze-generation analysis are described in Supplementary materials, “Eye tracking methods and generation attempt”.

### Participants

A total of fifty-eight healthy subjects with no significant mobility issues were recruited from community and a pool of university staff and students. Twenty-five subjects were recruited for experiment 1; one subject was excluded due to technical failure. The final sample included 24 subjects (17 female, mean age 25.71 years). Thirty-three subjects were recruited for experiment 2. Two subjects were excluded due to technical failure and compliance failure, respectively. The final sample included 31 subjects (15 female, mean age 24.81 years). An a priori power analysis was not conducted due to the novelty of the investigation and the complexity of the analyses. Instead, we based our target sample size (*N ≈* 30 per experiment) on previous studies using computational modelling of neurophysiological data (Mahajan et al., 2025), as well as EEG, SCR, and pain studies (Schulz et al., 2015; Zhang et al., 2018), and studies from our group using combined neurophysiological recordings and VR (Hewitt et al., 2026). This approach represents a pragmatic balance that ensures the credibility of the results and the stability of model estimates while accounting for the high per-subject cost and depth of data collected from each individual.

The procedure was approved by the Research Ethics Committee of the University of Oxford (CUREC approval reference R58778), and all participants gave written informed consent at the start of the experiment following the Declaration of Helsinki. Participants were reimbursed with £30-£45 depending on the duration of experiment, including a £10 performance incentive. Participants were informed at the start of the experiment that their total points would be rewarded with a monetary incentive of up to £10. To maintain a constant level of motivation throughout the task, the exact point-to-currency exchange rate was not specified. Upon completion of the session, all participants were awarded the maximum bonus of £10.

For analyses requiring eye-tracking data, four additional participants in each experiment were excluded if any analysed 60-s block contained more than 100 frames with an exactly identical `gaze direction` vector, counted across the block rather than requiring consecutive frames, leaving 20 participants in Experiment 1 and 27 in Experiment 2. An exploratory attempt to generate gaze from head movements using HAGI++ (Jiao et al., 2025a,b) is described in Supplementary materials, “Eye tracking methods and generation attempt”.

### Statistical Analysis

Statistical analysis was carried out using custom scripts written in Python 3 and R 4.4.2. For LMMs, lme4 and lmerTest packages were used (Bates et al., 2015; Kuznetsova et al., 2017). We initially fitted subject-specific intercepts and the random slopes specified for each analysis, allowing correlations among these random effects. When singular fits occurred within an analysis block (SCR models, EEG channel-wise fits or permutation batches), we used uncorrelated random effects for the entire block, including nonsingular fits, without removing any random intercepts or slopes. This limited simplification was chosen to support numerical stability and a reasonable balance between Type I error and power (Bates et al., 2018; Matuschek et al., 2017). Comparisons of maximal and no-correlation fits are provided in Supplementary materials, “Linear mixed model outputs”. Fixed effects were tested using *t*-tests with Satterthwaite-approximated degrees of freedom, as implemented in lmerTest. Confidence intervals for fixed effects are Wald intervals computed from the same quantities, i.e. the estimate ±*t*_1_*_−α/_*_2,_ *_ν_* times its standard error, where *ν* is the Satterthwaite degrees of freedom for that coefficient and *α* = 0.05. These intervals are therefore consistent with the reported *t* and *p* values by construction. Supplementary materials provide additional details of the formula of the LMMs and their detailed outputs.

### Model description and fitting

#### Experiment 1

As we disregarded planning, the model in experiment 1 compared *C_p_u*(*a*)+*C_m ·_ d*(*a*) for each fruit at a particular time point. The first question was what fruits were available for subjects to choose from. Fruits were abundant in the subjects’ reachable area, however, many were not directly in subjects’ visual field or hidden behind the trees. Subjects also needed to identify the fruits’ colour to know whether the fruit was painful or not. Therefore, there was an additional search (or exploration) process that was not modelled here. We utilised eye-tracking technology (HTC Vive Pro Eye), and assumed only fruits that were in the central visual field were considered by the subject. Fruits that were captured by eye-tracking were saved into a first-in-first-out memory queue (the available choice set). We heuristically set the memory size to be 7, and the memory was cleared each time the subject chose a fruit. Next question was when the subject made a decision. The first time point that the subject looked at a pineapple not seen before could be used as the decision point in this free-operant task. However, we could not evaluate the model when the chosen fruit was not in the memory yet, as the model would never make a correct prediction out of a set of wrong candidates. We could only retrospectively identify the fruit that the subject chose and only fitted the model when the correct choice was in the choice set. This acausal problem is difficult to solve without taking time into consideration. We will show in the experiment 2 model fitting, by quantifying the effort and opportunity costs as a function of time delay, the model-fitting process was made causal.

We had converted the problem into finding a set of hyperparameters so that at each decision point, the maximum value of all choice differential values *C_p_u*(*a*) + *C_m_* · *d*(*a*) predicts the choice that the subject choose. Although this deterministic argmax rule is related to the decision rule implied by multinomial logistic models, we did not specify a probabilistic softmax choice function or estimate the parameters by maximizing a multinomial likelihood. Instead, we selected the parameter values that minimized the unweighted mean of the block-specific misclassification rates, each calculated as the number of incorrectly predicted choices divided by the total number of choices in that block. The utility function *u* was parameterized as a sigmoid whose horizontal scale and location were also estimated. Because the resulting accuracy objective is discrete and need not be differentiable, we optimized it using an exhaustive grid search over the specified parameter values. The complete grid specification is provided in the Supplementary materials.

To provide a detailed description regarding the computational framework, we summarise the properties of the first optimality equation and the underlying state-action space in Table 2.

**Table 1.** Summary of statistical analyses, variables, and corresponding results.

| Result | Dependent variables | Independent variables | Model |
| --- | --- | --- | --- |
| Fig. 3A | Aversive choice probability | Pain ratings, Subject (Random slope & intercept) | LMM |
| Fig. 3B | Choice distance bias | Pain ratings, Subject (Random slope & intercept) | LMM |
| Fig. 3G | Phasic pain utility values | Pain ratings, Subject (Random slope & intercept) | LMM |
| Fig. 4B | Fixation SCR coefficients | Phasic pain utility values, Subject (Random slope & intercept) | LMM |
| Fig. 4B | Fixation SCR coefficients | Pain ratings, Subject (Random slope & intercept) | LMM |
| Fig. 4C | Shock SCR coefficients | Phasic pain utility values, Subject (Random slope & intercept) | LMM |
| Fig. 4C | Shock SCR coefficients | Pain ratings, Subject (Random slope & intercept) | LMM |
| Fig. 5A–B | Pain ratings | Phasic pain, Tonic pain, Subject (Random Factor) | 2-way RM ANOVA |
| Fig. 5C–D | Aversive choice probability | Pain ratings, Subject (Random slope & intercept) | LMM |
| Fig. 5E | Aversive choice probability | Phasic pain, Tonic pain, Subject (Random Factor) | 2-way RM ANOVA |
| Fig. 6 | N1–P2 amplitude | Phasic pain, Tonic pain, Subject (Random Factor) | 2-way RM ANOVA |
| Fig. 9B | Vigour constant $C_v$ | Phasic pain, Tonic pain, Subject (Random Factor) | 2-way RM ANOVA |
| Fig. 9E | Phasic pain utility values | Aversive choice probability | Linear regression |
| Fig. 10A | Band power | Tonic pain, Head speed, Subject (Random slope & intercept) | LMM |
| Fig. 10B | Band power | Vigour constant ($C_v$), Head speed, Subject (Random slope & intercept) | LMM |

**Table 2.**
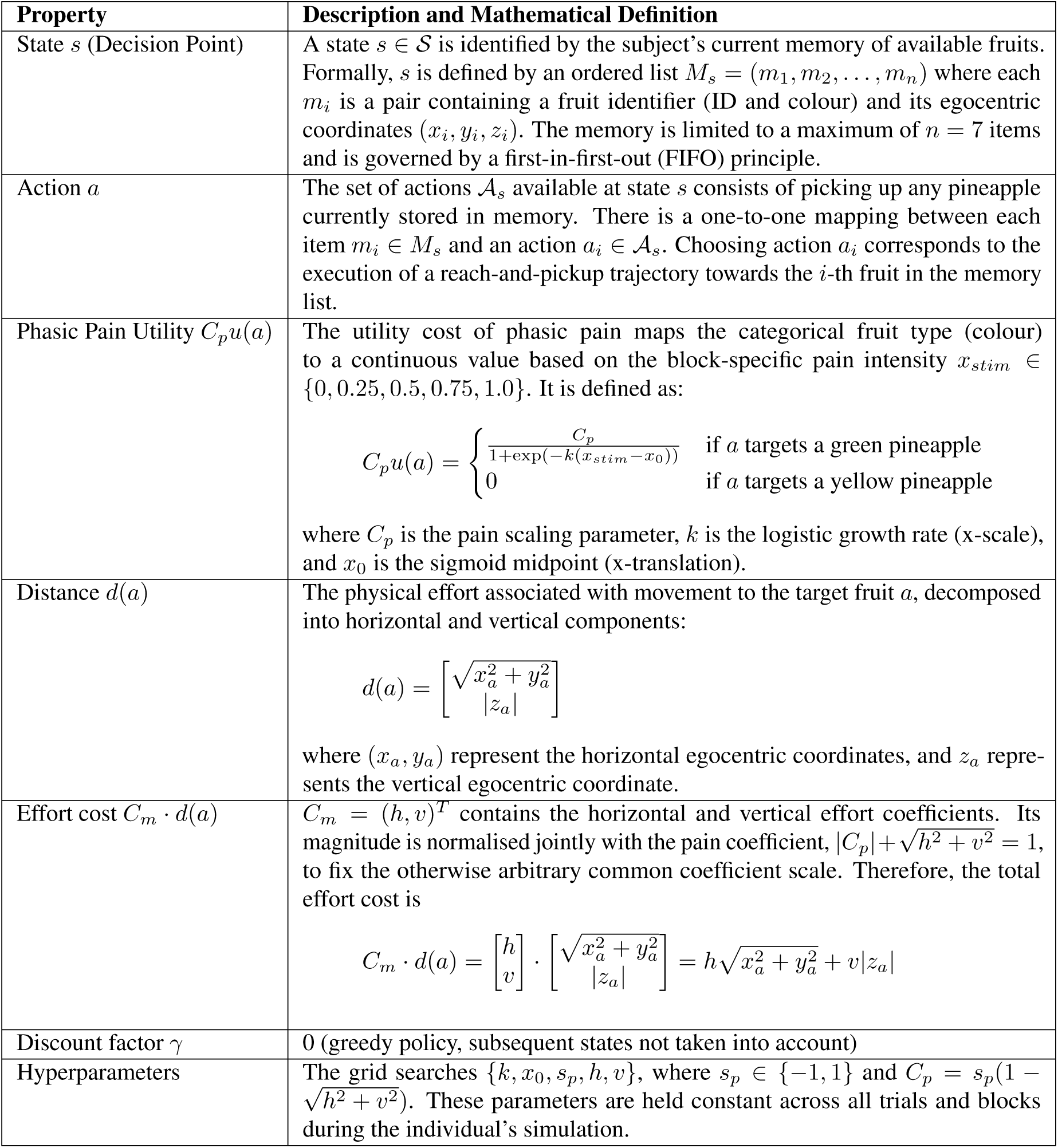
Summary of model properties for the Experiment 1 foraging task.

The simulation and model-fitting process are conducted blockwise. Because the phasic pain intensity *x_stim_* for green fruits is fixed for the duration of a single block, this information is treated as a known variable to the model rather than being explicitly encoded into the state space *s*. A discussion on the choice of constant *x_stim_* for the duration of a single block is added in the Supplementary materials section: Discussion of Pain Intensity Information and Model Robustness.

Furthermore, we clarify the distinction between parameters and hyperparameters in this framework. Unlike traditional reinforcement learning models where internal parameters (such as action values or learning rates) may be continuously updated through trial-and-error, we have eschewed learning to maintain simplicity and focus on the steady-state motivational trade-offs. Consequently, there are no “online” parameters; the hyperparameters listed in Table 2 are the only variables optimised via the fitting procedure to describe the subject’s stable behavioural preferences across the experimental conditions.

#### Experiment 2

A key assumption in Niv et al. (2007) was that animals chose an action pair of an action and a delay. The delay included both the time waiting and the time executing the action. It was an approximation of real animal behaviour, and in fitting to real human subjects data in this task, we did not have direct access to subjects’ delay decision. Therefore, we tracked the delay and computed the differential value at each time point. The central assumption in this model fitting solution was all differential values for the choices that were not chosen should be less than the value for the final chosen choice. The model-fitting program tracked the model predicted value for each fruit that a subject fixated on. The model’s predicted choice was correct (i.e., fit the data) when the chosen fruit had the highest predicted value (compared to other fruits within the time interval before the last fixation on the chosen fruit). Otherwise, the model’s predicted choice was incorrect. *γ* is still set to 0 for model-fitting in this experiment.

**Table 3.**
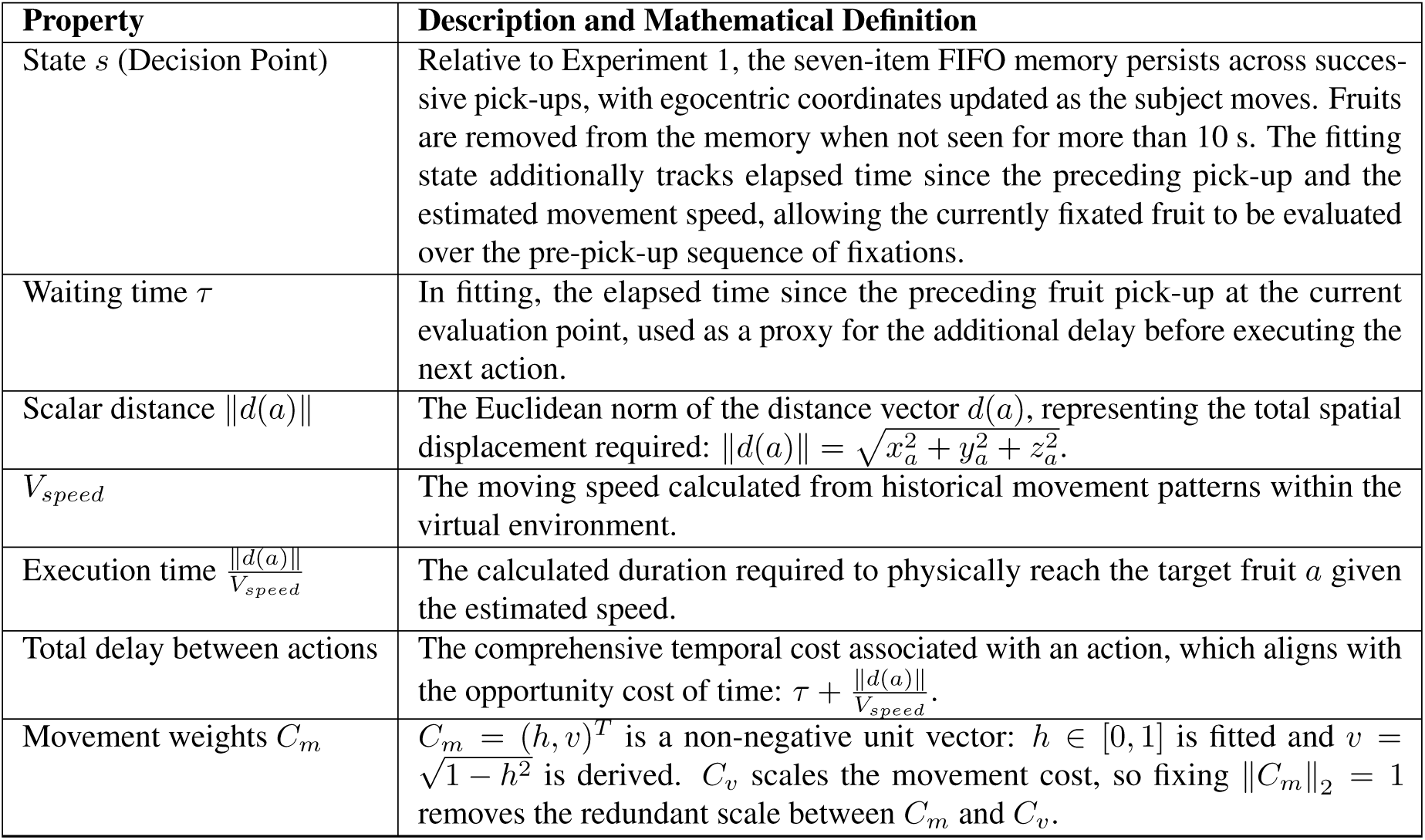
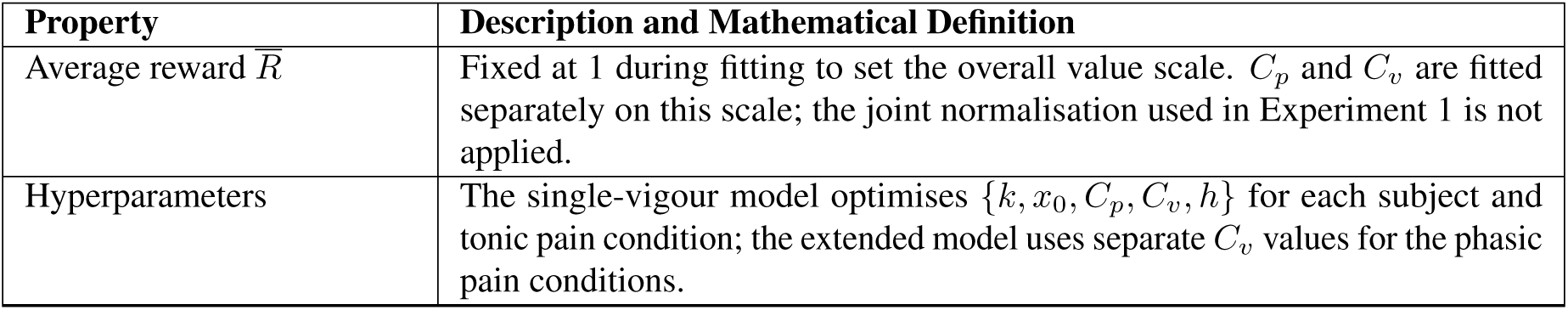
Summary of model properties for the Experiment 2 foraging task (addendum to Experiment 1 Table)

**Figure 11.**
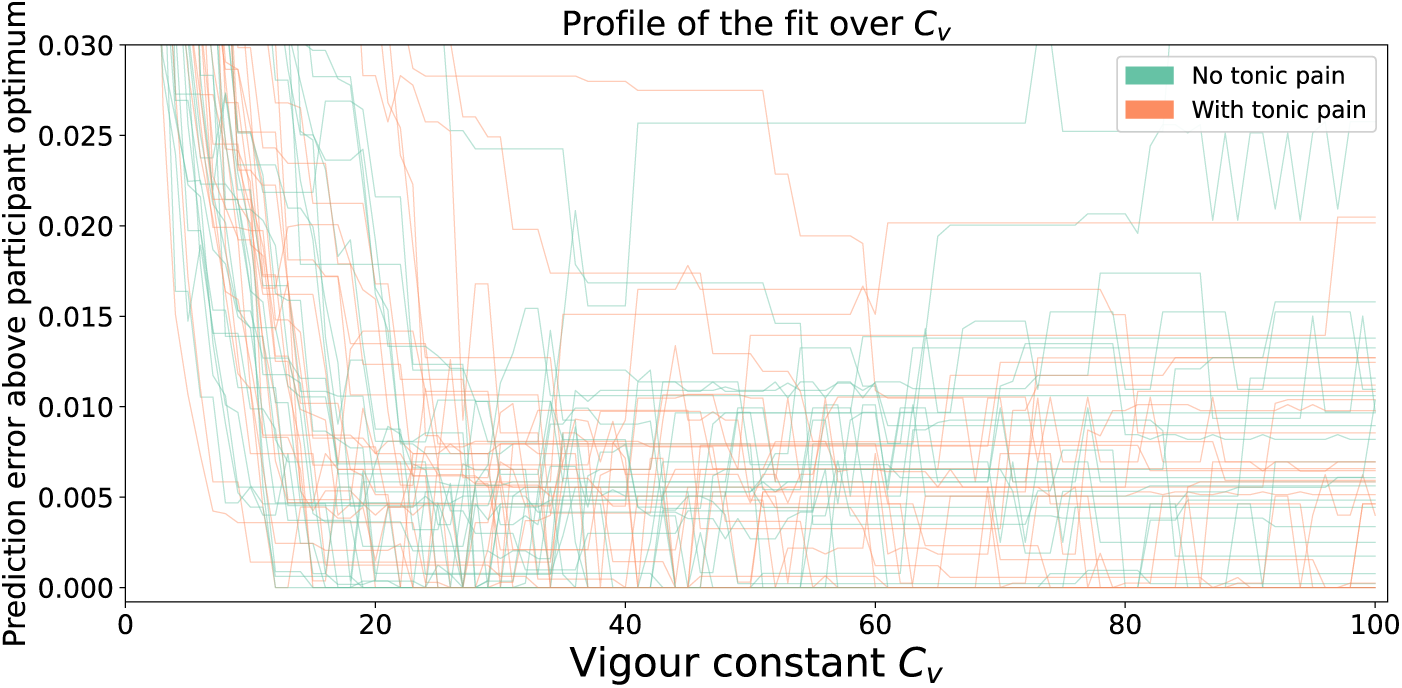
Profile of the fit over *C_v_* in the single-vigour model. Each curve represents one participant and tonic condition (*n* = 27 per condition). At each integer *C_v_* from 1 to 100, all other fitted parameters were reoptimised over the grid. The horizontal axis shows *C_v_*; the vertical axis shows excess block-averaged prediction error relative to that participant’s minimum in the same condition. Lines join the sampled values. Only excess errors up to 0.03 are displayed. These are error profiles, not confidence intervals, and the upper extent of optimal sets reaching 100 is unresolved within the grid. Note the existence of non-global local minima on the profile lines.

#### Vigour constant fitting sensitivity

To assess parameter precision, we profiled the single-vigour fit over *C_v_* = 1*,…,* 100. At each integer value, the profile records the lowest block-averaged choice-prediction error obtainable by reoptimising all other parameters on the fitting grid (Fig. 11). The original ordered grid search retains the first optimal joint parameter tuple, which need not contain the smallest profiled *C_v_*.

The exact optimal sets were broad: their median spans (maximum minus minimum optimal *C_v_*) were 13 units with tonic pain and 11 without tonic pain. Across selections of exact optima for each participant and condition, the mean tonic-minus-no-tonic *C_v_*contrast had attainable extremes of 4.48 and +31.74, compared with +15.59 under the original selection convention. Each extreme permits the same minimum error in every participant and condition, with the other parameters reoptimised. These are sensitivity bounds within the searched grid, not confidence limits; they depend on the upper boundary of 100 and do not imply that every intermediate contrast is attainable. The reported positive contrast and its statistical significance are therefore sensitive to the selection among equally fitting solutions.

Choosing the nearest considered fruit offers an intuitive explanation for weak upper bounds on *C_v_*. Increasing *C_v_* can make further alternatives less attractive, so the choice may require a minimum value of *C_v_* without imposing an upper limit. An upper bound would instead require a rejected alternative in the candidate set that becomes preferable as *C_v_* increases, such as a closer fruit, which occurred infrequently in the candidate sets. Equally good fits may consequently extend beyond 100. This gives a practical rationale for retaining the first minimum encountered in the ascending joint-grid search as a reproducible representative. It does not resolve the non-identifiability of optimal *C_v_* or establish a unique optimum.

### Experimental pain stimulation

#### Electric phasic pain

The electric stimulation pulses were generated by a DS5 isolated bipolar constant current stimulator (Digitimer, Letchworth Garden City, UK) and delivered using a Wasp pain stimulation electrode. These electrodes preferentially activate nociceptive A-delta fibres, thereby eliciting ERPs that more accurately reflect nociceptive processing compared to standard bipolar stimulation (Inui et al., 2002; Mørch et al., 2011). The pulses were 200 Hz square waves with a 1 ms pulse width.

Calibration of stimulation intensity was conducted based on subjects’ pain ratings before the task commenced. To maintain consistency in pain ratings, additional calibration could be performed at the end of the 10th block in experiment 1 or at the end of every six blocks in experiment 2. Subjects rated their pain on a 0-10 numerical rating scale while researchers adjusted the stimulation intensities following a staircase procedure. The calibration process was considered complete when subjects provided consistent pain ratings within the desired range, as determined by the experimental design (8/10 in experiment 1 for the max pain condition; 3/10 in experiment 2 for the low phasic pain condition and 7/10 for the high phasic pain condition). At the end of each block, subjects provided a pain rating for that block. Ratings in both procedures were given verbally. During calibration, researchers recorded the subjects’ verbal ratings, whereas in post-block ratings, verbal responses were processed using speech recognition services (Microsoft, Redmond, US) and automatically converted into a visual analogue scale bar displayed in VR. In case of speech recognition failure, researchers wrote down the ratings and manually updated the data for analysis.

#### Pressure tonic pain

We used pressure cuffs (VBM, Sulz, Germany) to safely induce tonic pain (Graven-Nielsen et al., 2015; Joseph et al., 2022). The cuff was wrapped around the non-dominant arm’s biceps. The cuffs were inflated prior to the start of each required block and immediately deflated after the block ended. Subjects were asked to report verbally when they felt discomfort and when they felt pain as the researcher manually inflated the cuff. Inflation stopped once the subject reported pain. To control for variables associated with wearing the cuff other than pain, pressure cuffs remained wrapped around the subjects’ arms even in blocks without tonic pain.

### Skin conductance

#### Data collection and processing

Experiment 1 was designed to establish the robust behavioural effects of the foraging task while ensuring the collection of reliable physiological data. We chose SCR as it is a well-validated index of autonomic arousal that we were confident would provide a clear peripheral measure of pain-related processing in this novel VR paradigm. In Experiment 1, SCRs were collected using a custom-built wireless sensor. The microcontroller was an Arduino Nano 33 IoT (Arduino SA, Chiasso, Switzerland). The skin conductance sensor was a Grove GSR sensor (Seeed Studio, Shenzhen, China). The firmware was written by the authors of this article with a designated sampling rate of 1000 Hz.

Data was segmented by experimental blocks (60 seconds long each). An entire block of data was removed from further analysis if the data loss was more than 20% of the expected total number of samples due to temporary wireless transmission failure. Each block of data was down-sampled to 10 Hz, and re-centred by the average baseline. A median filter with a window size of 5 was applied to the down-sampled data to further remove artefacts. Further small gaps of missing data were handled in the ordinary least square fitting in R with `na.omit` as the default option. As subjects would only know the intensities of pain stimulation in new blocks after picking up a painful fruit, for fixation events, only events after picking up the first painful fruit were analysed. For the wireless recordings, we improved clock synchronisation with task events by applying a block-specific affine correction for clock offset and drift to the Arduino timestamps (Supplementary materials, Clock synchronisation).

### Effect coefficient estimate

Linear regression were used to fit the skin conductance time series data to the constructed time series data. The time series was generated by convolving the event trigger with the CRF, with hyperparameters taken from (Bach et al., 2010). Fixation events that occurred during picking up painful fruit were not included as shock-evoked SCR was strong. For computational practicality, convolutions were truncated between 0 and 60 seconds (the length of an experimental block) (Bach et al., 2009). Each linear regression was fitted separately to individual blocks, generating its own set of coefficient estimates.

Block-level coefficients were retained irrespective of their first-level *p* values. For each event type, coefficients with absolute *z*-scores of 3 or greater across the pooled blocks were excluded. The remaining coefficients, including negative estimates, were analysed using unweighted linear mixed-effects models with SCR coefficient as the outcome and either decision value or subjective rating as the predictor. Predictors were standardised across the experiment’s block-level data before outcome-specific filtering. We also tested inverse-variance mixed-effects models using `metafor` and `brms`, incorporating heteroskedasticity- and autocorrelation-consistent (HAC) estimates of the block coefficients’ sampling variances and allowing for additional block-level heterogeneity. These analyses were applied to both experiments; their specification and results are reported in Supplementary materials, Skin conductance sensitivity analyses.

### EEG

#### Data collection and processing

Whole-scalp EEG data was continuously recorded with 32-channel LiveAmp system (BrainProducts GmbH, Munich, Germany) at 500Hz. We chose FCz as the reference. To remove ocular artefacts, recordings were first cleaned with EEGLAB (v2023.1) by removing channel drift (linear filter (FIR) transition band from 0.25 to 0.75 Hz). It also automatically removes bad data periods by thresholding max acceptable 0.5 second window standard deviation of 20, max acceptable channel RMS range of 7, and maximum out-of-bound channels of 25%. The ICA was then performed on the cleaned data and the weight matrix was computed and saved separately. Ocular artefact components were identified manually, assisted by automated classifier ICLabel (Pion-Tonachini et al., 2019). We then cleaned ocular artefacts by removing these identified components back projected onto the raw data. By manually inspecting the data, we interpolated up to 10% (3 channels) of electrodes. The output data at this stage served as the input data for ERP and time-frequency analysis separately. For analyses aligned to timestamped task events recorded on the PC, we corrected relative clock offset and drift using a recording-specific affine mapping from PC timestamps to EEG sample indices, estimated from paired trigger records (Supplementary materials, Clock synchronisation).

#### ERP analysis

ERP analysis was conducted on the preprocessed data in Python with MNE-Python 1.7.0 package. The data was re-referenced to the average amplitude of all electrodes. A bandpass filter from 1 to 30 Hz was applied. The high-pass cutoff frequency was chosen because of the presence of strong motion artefacts in our mobile EEG data. Identical analysis was applied at 0.1Hz and 0.5Hz high-pass cutoff frequencies, and significant results remained unchanged (see Supplementary figures). Recordings were processed into 1s epoch with 0.3s baseline before the event trigger. Epochs with peak-to-peak amplitude exceeding 200 *µ*V in Cz channel were excluded (average rate of excluded epochs was 2.81%). We searched for N1 amplitude peaks from 100-170ms and P2 amplitude peaks from 140-300ms at channel Cz (Favero et al., 2023; van den Broeke et al., 2010).

#### LMM-based time-frequency analysis

Time-frequency analysis was performed on preprocessed data in Python with custom scripts relying on MNE-Python 1.7.0 package. The data was re-referenced to the average amplitude of all electrodes. A notch filter with 4Hz width centred at 50Hz was applied. Data were epoched from 0-0.5s according to the decision points. The decision points were chosen when subjects fixated on the chosen fruit for the first time, or at a re-evaluation point. A re-evaluation point was a subsequent fixation on the chosen fruit following fixation on another available fruit or collection of the model’s stored best-valued alternative, irrespective of whether the chosen fruit’s value exceeded competing values.

The spectral power was estimated using Welch’s method. A Hamming-tapered window was used with the window length equal to the epoch length (250 samples, 0.5 second). The estimated spectral power was separated into four bands: theta 4-7 Hz, alpha 8-12 Hz, beta 13-29 Hz, gamma 30-100 Hz (Levy et al., 2023; Chrastil et al., 2022; Schulz et al., 2015), and averaged within each band range. These band-averaged values formed the basis for the LMM-based time-frequency analysis presented in the main text, which was designed to statistically account for the task’s complex noise profile. For each power band, averaged band power was inputted into an LMM for each channel as the dependent variable (Schulz et al., 2015). For each epoch, the z-score of the averaged band power was computed for each channel within the subject, and the epoch would be removed from LMM fitting for this band if any channel had a z-score greater than 3 (Nolan et al., 2010). For tonic pain effect, the binary condition of tonic pain presence was used as an independent variable. Random effects for tonic pain and intercepts were estimated without an intercept–slope correlation for the topographies reported here. Crucially, a single head movement speed variable was added as an additional independent variable to reduce motion artefacts. The head movement speed for a particular epoch was calculated by averaging the speed between 10 Unity frames before the decision point and 10 Unity frames after the decision point. For vigour constants, the LMM had the same structure except replacing the binary tonic pain condition by fitted vigour constants presented in Figure 9A. Source analysis for spectral power was included in the Supplementary materials (section: Source Analysis). Complementary analyses of total event-related and ERP-subtracted induced responses to phasic pain, including tonic-by-phasic intensity interaction tests, are provided in the Supplementary materials (section: Event-related time–frequency and induced oscillatory responses to phasic pain stimuli).

#### Cluster-based permutation analysis

We additionally assessed the spatial extent and strength of the EEG associations using cluster-based permutation tests (Maris and Oostenveld, 2007). Tests were performed separately for tonic pain condition and the condition-specific fitted vigour constants, within each frequency band. The same post-exclusion epoch data and LMM formula were used for the observed and permuted maps: band power was predicted by the variable of interest and head-movement speed, with subject-specific random intercepts and slopes for the variable of interest and no intercept–slope correlation. At each electrode, the fixed-effect *t* statistic and its Satterthwaite-approximated degrees of freedom were obtained using lmerTest. Each *t* statistic was converted to a signed standard-normal deviate with the same tail probability. Spatially adjacent electrodes exceeding |*z >*| 1.96 (two-sided uncorrected *p <* .05) were grouped into separate positive and negative clusters. Electrode adjacency was derived by Delaunay triangulation of the sensor positions using MNE-Python. Cluster mass was the sum of the *z* values within each cluster.

For each observed map containing a suprathreshold cluster, 1,000 permutations shuffled the block-level predictor values within each participant and each of the three consecutive six-block groups, preserving the number of blocks assigned each value. For vigour, the two condition-specific fitted constants for each participant were held fixed and reassigned among blocks. A permuted block value was applied to every epoch and electrode in that block, while EEG observations and head-movement values remained attached to their original epochs. All electrode-wise LMMs were refitted and clusters recomputed for each permutation. The largest absolute cluster mass across both signs was retained, with zero assigned when no cluster formed. For each observed cluster, the permutation *p* value was calculated as (*b* + 1)*/*(1000 + 1), where *b* was the number of permuted maxima at least as large as the absolute observed cluster mass (Phipson and Smyth, 2010). Cluster *p* values below .05 were considered significant. This correction was applied across spatial clusters within each frequency band and predictor, without further correction across bands or predictors. Because only the predictor was shuffled, the permutations did not preserve its observed dependence on head movement; this limits interpretation of the test as an exact test of the association conditional on movement (Winkler et al., 2014).

## ACKNOWLEDGMENTS

ST would like to thank Pranav Mahajan, Yijia Yan, Suyi Zhang, and Eoin Kelleher for their feedback on the content of this paper during the work-in-progress stage. ST also thanks Kirsty Bannister, Joseph Taylor, Kristian Hennings for sharing valuable knowledge on the safe use of pressure cuffs to induce tonic pain. Special thanks to Eoin Kelleher for providing an exceptional visual representation of the task (Fig. 1A). ST would like to thank the reviewers of IASP 2022, CCN 2023, and COSYNE 2024 for their feedback.

The work was funded by Wellcome Trust (214251/Z/18/Z, 203139/Z/16/Z and 203139/A/16/Z), IITP (MSIT 2019-0-01371, RS-2023-00233251), JSPS (22H04998), and EPSRC (EP/W03509X/1). This work was also funded by the EPSRC and MRC Research Grant Scheme under the reference number UKRI1970. This research was also partly supported by the NIHR Oxford Health Biomedical Research Centre (NIHR203316) and the Royal Academy of Engineering. The views expressed are those of the author(s) and not necessarily those of the NIHR or the Department of Health and Social Care. For the purpose of open access, the author has applied a CC BY public copyright licence to any Author Accepted Manuscript version arising from this submission.

## DATA AND CODE AVAILABILITY

The multimodal behavioural and physiological dataset has been de-identified. Access to the full dataset is restricted as an additional precaution to safeguard participant privacy, given the detailed and linked nature of the data. Researchers seeking access should contact the corresponding authors, Shuangyi Tong or Ben Seymour. Requests will be considered on a case-by-case basis in accordance with applicable University of Oxford information governance and data protection policies, relevant ethical requirements and appropriate data-sharing terms.

The complete analysis code is publicly available at https://github.com/ShuangyiTong/Phasic-and-tonic-pain-serve-distinct-functions-during-adaptive-behaviour. A sample dataset has been released in the same repository to illustrate the custom data structure and support exploration of the analysis code.

Software tools for replicating the experiment are available at https://github.com/ShuangyiTong/PineappleStudy2025ReplicationSoftware.

## AUTHOR CONTRIBUTIONS

ST and BS designed the theory and experimental protocols. ST, BS, and TD developed the VR research platform used for the experiments. ST collected and analysed the data. ST and DH designed the EEG analysis pipeline and methodology. SL and TD provided critical feedback and theoretical insights throughout multiple iterations of the study. ST and BS wrote the first draft. ST, TD, DH, SL, and BS edited and approved the manuscript. TD, SL, and BS acquired funding for the study.

