## Supplementary Materials for "Phasic and tonic pain serve distinct functions during adaptive behaviour"

#### ABSTRACT

The supplementary materials for the paper "Phasic and tonic pain serve distinct functions during adaptive behaviour" are presented below.

#### CONTENTS

**1 Source Analysis 2**
**2 ERP with different band pass filter 2**
**3 Event-related time–frequency and induced oscillatory responses to phasic pain stimuli 3**
**4 Linear mixed model outputs 8**
**5 Possible numerical differences in large LMM fitting across different hardware platforms 9**
**6 Discussion of Pain Intensity Information and Model Robustness 9**
**8 Skin conductance sensitivity analyses 12**
**9 Grid search parameter configuration 13**
**10 Eye tracking methods and generation attempt 15**
**References 17**

### 1 SOURCE ANALYSIS

In this section, we presented the results of the source analysis, which supplemented the channel-wise topography plots. Following the pre-processing described in the Methods section, band-pass filters were applied to the pre-processed signals and epoched for the same 0–0.5 s after decision points as in the time-frequency analysis. Epochs rejected in time-frequency analysis were rejected for source analysis. The leadfield matrix was computed for a recursively subdivided icosahedron brain surface, consisting of a total of 1,284 vertices derived from MNE-provided templates. The sLORETA method was then applied to each band-passed signal to obtain a time-domain signal for each source location (Pascual-Marqui, 2002b). For each source location, the Welch method was used to estimate the total spectral power of each epoch. The same LMM employed in the time-frequency analysis was applied, except that the band power in the time-frequency analysis was replaced by the total spectral power estimated from inverse solutions of band-passed signals at source locations (Frei et al., 2001; Gil Ávila et al., 2023; Pascual-Marqui, 2002a).

Figure S1 shows the t values from the LMM's output. Consistent with the channel-wise topography results, we observed decreased signal power for Alpha bandpassed signal in the right hemisphere. Specifically, most noticeable negative correlation with tonic pain showed up in parietal regions for Alpha bandpassed signal.

### 2 ERP WITH DIFFERENT BAND PASS FILTER

In the main text, the ERP was processed with a bandpass filter from 1-30 Hz. We present additional 0.5-30 Hz and 0.1-30 Hz bandpass filtered ERP results here.

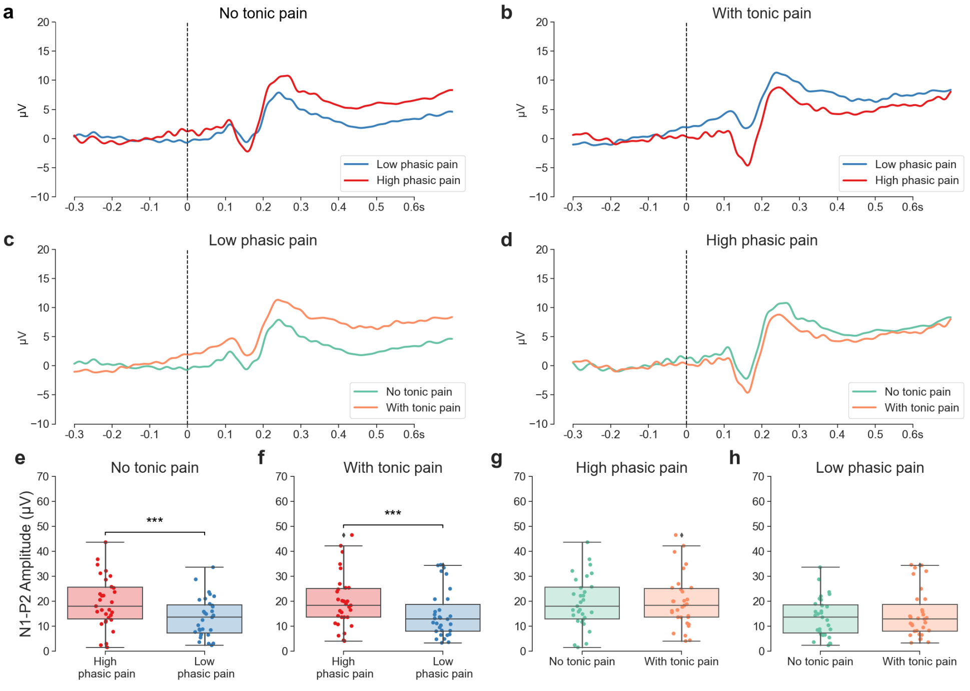

**Figure S2.** Phasic pain ERP at Cz, band-passed by 0.5-30 Hz filter. Two-way repeated measures ANOVA showed the effect for phasic pain was significant, $F(1, 30) = 33.58$, $p < .001$. The effect for tonic pain was not significant, $F(1, 30) = 1.21$, $p = .280$, and the interaction effect was also not significant, $F(1, 30) = 0.49$, $p = .489$

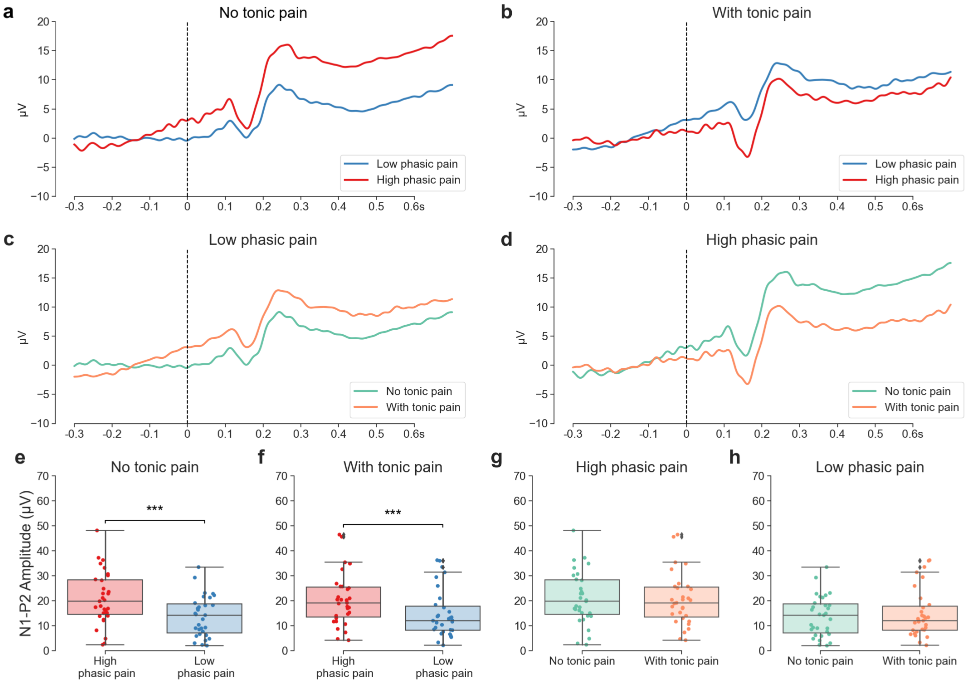

**Figure S3.** Phasic pain ERP at Cz, band-passed by 0.1-30 Hz filter. Two-way repeated measures ANOVA showed the effect for phasic pain was significant, $F(1, 30) = 47.60$, $p < .001$. The effect for tonic pain was not significant, $F(1, 30) = 0.26$, $p = .611$, and the interaction effect was also not significant, $F(1, 30) = 1.34$, $p = .256$

### 3 EVENT-RELATED TIME–FREQUENCY AND INDUCED OSCILLATORY RESPONSES TO PHASIC PAIN STIMULI

We examined two complementary time–frequency representations (TFRs) of the EEG response to phasic pain at electrode Cz: total event-related power, which includes both evoked and induced contributions, and induced power after removal of the condition-specific average evoked response. Both analyses used the same retained epochs from 31 participants.

##### *Preprocessing and Epoching*

Following the same preprocessing procedure described in the main text, the EEG signal was re-referenced to the common average, high-pass filtered at 1 Hz, and a notch filter at 50 Hz was applied to remove power line noise. Epochs were extracted from $-0.55$ s to $0.95$ s relative to stimulus onset. Trials with peak-to-peak amplitude exceeding $200\,\mu$V at Cz were excluded.

Retained epoch counts were limited and unequal across participants and conditions (Table S1). Thirteen of the 31 participants contributed fewer than 10 epochs in at least one analysed condition, and five contributed fewer than five. The same retained epochs were used to estimate each participant's condition-specific ERP and to average residual single-trial power after ERP subtraction.

**Table S1.** Retained epochs used in the phasic pain TFR analyses. Statistics describe the distribution of epoch counts across 31 participants within each condition. SD is the sample standard deviation, calculated with denominator $n-1$, where $n=31$ participants.

| Phasic pain | Tonic pain | Minimum | Maximum | Mean | SD |
|---|---|---|---|---|---|
| High | Absent | 3 | 49 | 15.97 | 11.32 |
| Low | Absent | 3 | 39 | 23.32 | 7.04 |
| High | Present | 3 | 36 | 15.00 | 8.77 |
| Low | Present | 3 | 41 | 23.90 | 9.29 |

##### *Time-Frequency Decomposition*

We used Morlet wavelet convolution at 40 geometrically spaced frequencies from 4 to 100 Hz, with the number of cycles set to $f/2$ for frequency $f$ in Hz. For total event-related power, we computed power separately for each epoch and then averaged power within each participant and condition. For induced power, we first subtracted each participant's condition-specific ERP from every epoch in that condition, and then computed and averaged the power of the residual signals. Thus, averaging was performed after power estimation in both analyses; only the induced analysis removed the average phase-locked response before decomposition.

For each representation, participant and condition, the trial-averaged power was expressed as $\log_{10}(P/P_{\mathrm{baseline}})$, where $P_{\mathrm{baseline}}$ was the mean power from $-0.3$ to 0 s at each frequency in that same representation. TFRs were cropped to $-0.3$ to $0.7$ s for plotting and statistical analysis. Grand averages were calculated across participants after this baseline correction.

##### *Statistical Analysis*

For each power representation, we tested four simple within-participant contrasts: high minus low phasic pain separately without and with tonic pain, and no tonic pain minus tonic pain separately at high and low phasic pain. We additionally tested the tonic-by-phasic intensity interaction by comparing each participant's high-minus-low difference with tonic pain against their high-minus-low difference without tonic pain. All five participant-level contrast maps were tested separately using one-sample cluster-based permutation $t$-tests (10,000 participant-wise sign-flip permutations, two-tailed) on the baseline-normalised log-power scale. The cluster-forming threshold corresponded to an uncorrected two-tailed $p < .05$; cluster-level significance was assessed at $p \leq .05$ against the permutation distribution of the maximum absolute cluster mass. This procedure controls multiple comparisons across the time–frequency plane within each contrast. No additional correction was applied across the five contrasts or the two power representations, so these supplementary analyses are exploratory.

##### *Results*

**Total event-related power.** The grand-average TFRs (Figure S4) showed a prominent post-stimulus increase in low-frequency power across all four conditions. High minus low phasic pain yielded one significant positive cluster without tonic pain ($p = .0001$) and one with tonic pain ($p = .0022$; Figure S5). Neither direct comparison between tonic conditions reached significance: the smallest cluster-level $p$ values were .0545 at high phasic intensity and .3163 at low phasic intensity. These results describe total event-related power, including evoked contributions.

**Induced power.** After subtraction of the condition-specific ERP (Figure S6), the high-minus-low phasic pain contrast retained one significant positive cluster without tonic pain ($p = .0001$), whereas no cluster reached significance with tonic pain (smallest cluster-level $p = .1925$; Figure S7). Neither direct tonic-condition contrast reached significance (smallest cluster-level $p = .0532$ at high phasic intensity and .5116 at low phasic intensity).

The direct tonic-by-phasic intensity interaction test identified no significant clusters in either total event-related power (smallest cluster-level $p = .8382$) or induced power (smallest cluster-level $p = .1575$). Thus, the different significance outcomes for the two high–low induced-power contrasts were not accompanied by a significant interaction on the analysed log-power scale. These null findings do not establish equivalence of neural responses across tonic conditions or exclude other tonic influences on phasic pain processing.

Interpretation is further limited by the small and unequal trial counts, particularly for high phasic pain. The free-operant task allowed participants to avoid painful outcomes rather than requiring a fixed number of high-pain trials. Across the 62 participant-by-tonic-condition counts from 31 participants, the number of retained high-pain epochs ranged from 3 to 49 (mean 15.48, sample SD 10.06). With few trials, non-phase-locked activity and noise may not cancel in the estimated ERP, so subtracting that estimate can also remove some of these components from the individual trials. In addition, trial-to-trial variation in evoked amplitude or latency can remain in the residual signal after average-ERP subtraction. These limitations reduce the precision of the estimates and the separation of evoked and induced activity; they warrant caution without establishing that the observed cluster is an artefact.

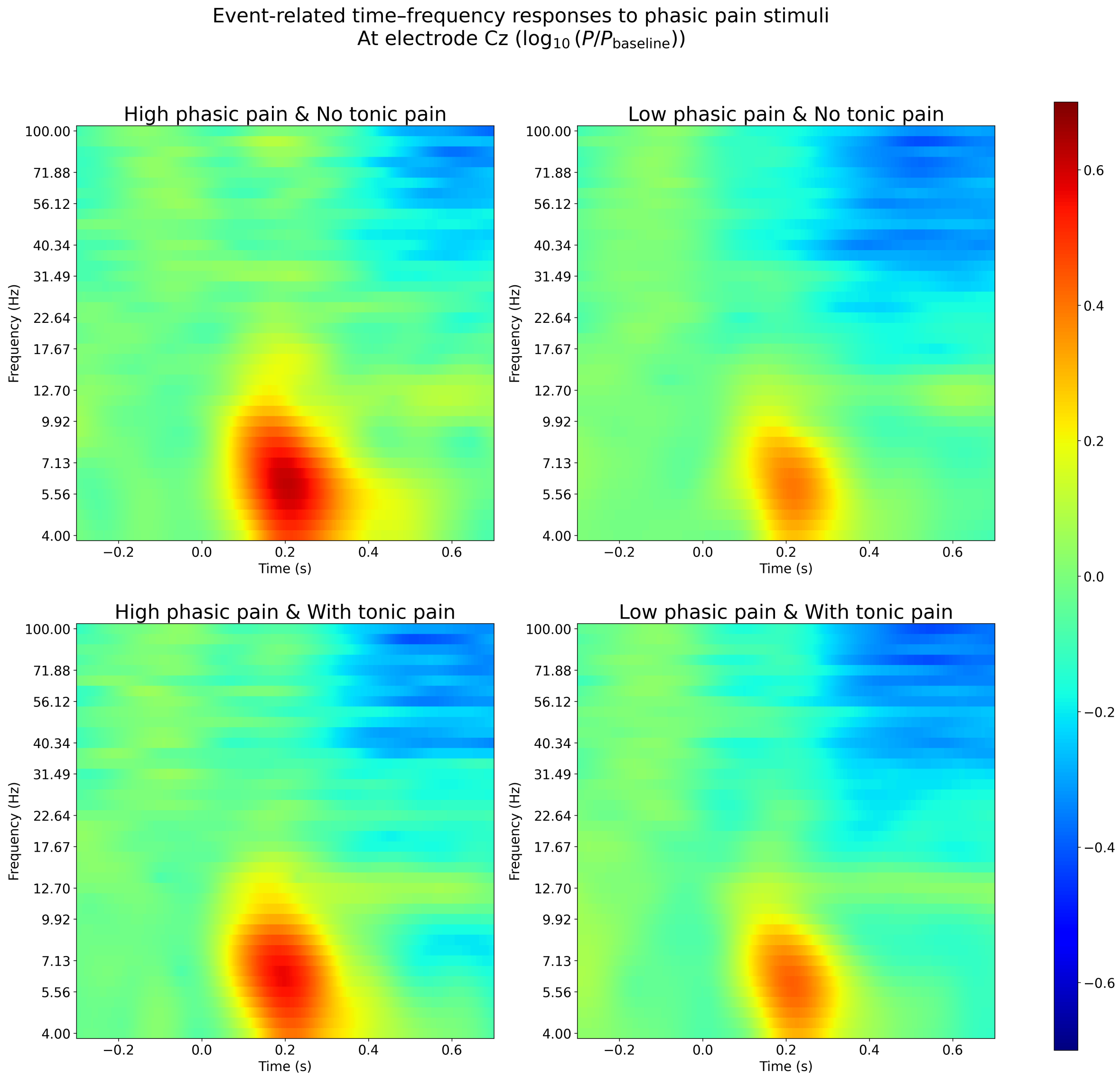

**Figure S4.** Grand-average event-related time–frequency responses to phasic pain at Cz. Total power was calculated from individual epochs before averaging, retaining both evoked and induced contributions. Panels show the four combinations of phasic intensity and tonic condition. Colours represent $\log_{10}(P/P_{\text{baseline}})$ relative to the $-0.3$ to $0$ s baseline. Time zero denotes phasic stimulus onset.

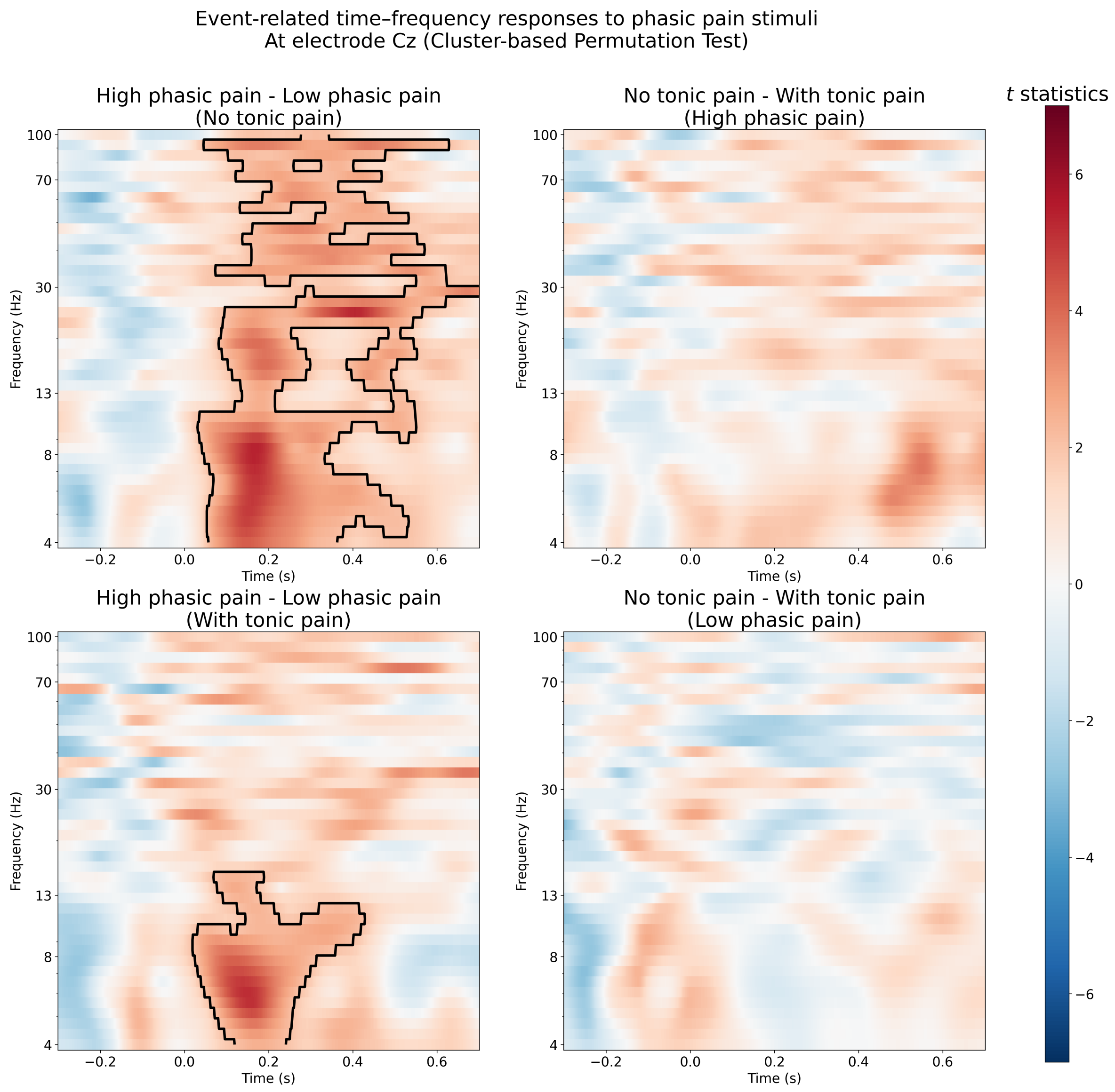

**Figure S5.** Statistical contrasts of total event-related power at Cz. Maps show $t$ statistics for the four within-participant contrasts. Black contours indicate clusters with permutation-corrected $p \leq .05$ within each contrast. High–low phasic pain differences were significant both without tonic pain (one cluster, $p = .0001$) and with tonic pain (one cluster, $p = .0022$); no significant clusters were observed for the direct tonic-condition contrasts. Correction was performed separately for each contrast and power representation.

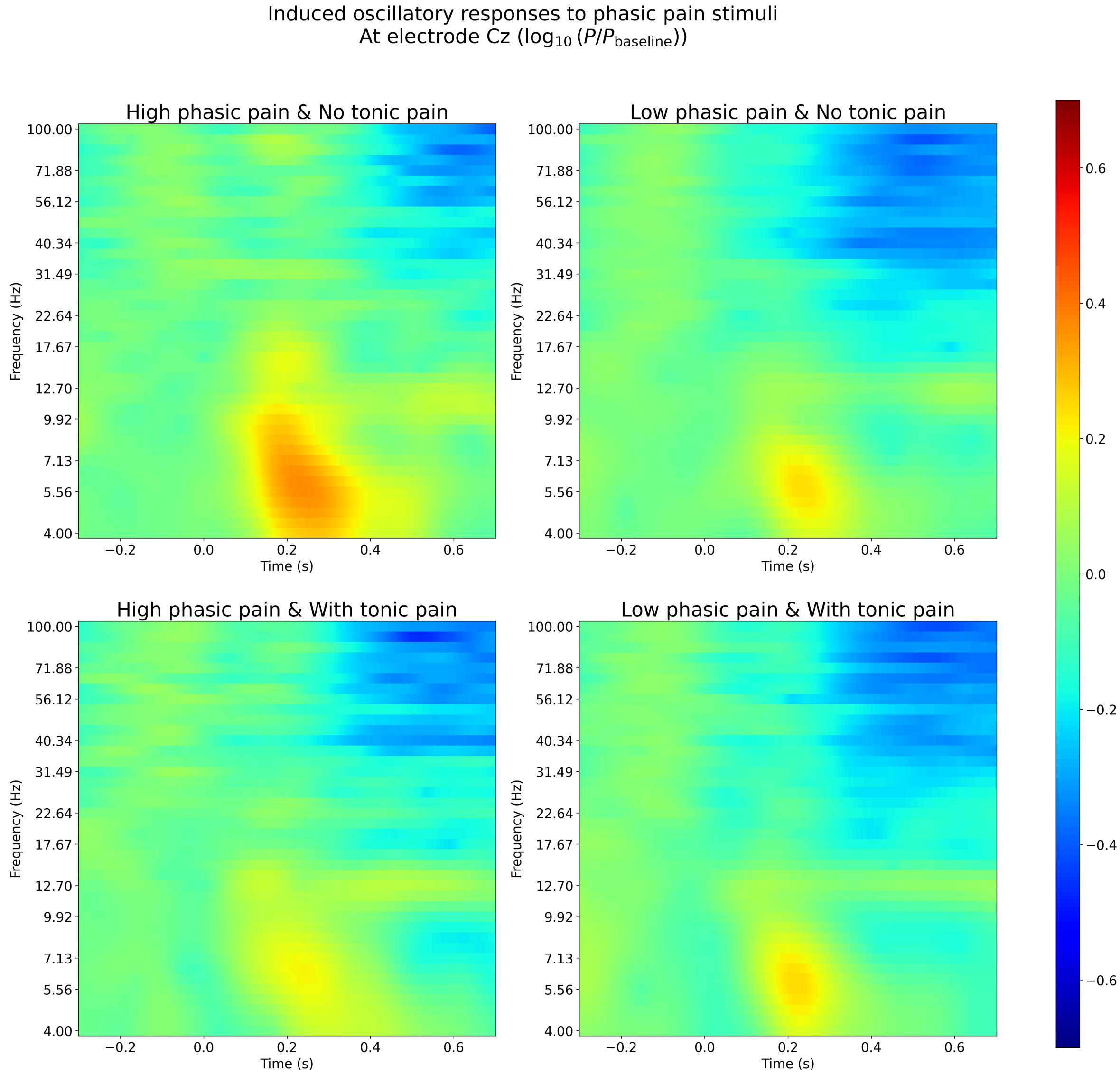

**Figure S6.** Grand-average induced oscillatory responses to phasic pain at Cz. Each participant's condition-specific ERP was subtracted from the individual epochs before calculating and averaging power. Panels and colour limits match Figure S4; baseline correction uses the induced-power baseline for each participant and condition.

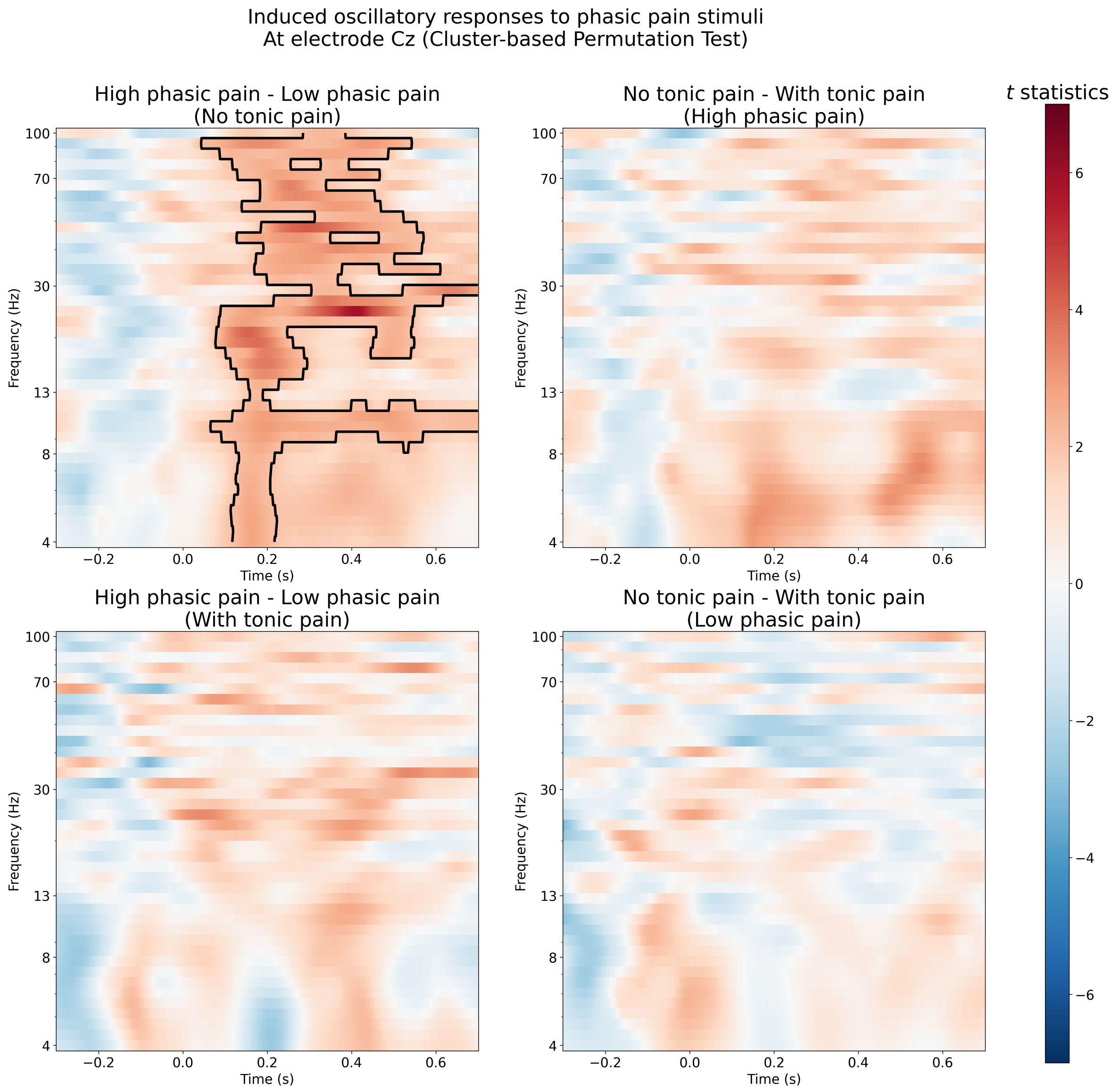

**Figure S7.** Statistical contrasts of induced power at Cz. Black contours indicate clusters with permutation-corrected $p \leq .05$ within each contrast. The high–low phasic pain contrast reached significance without tonic pain (one cluster, $p = .0001$) but not with tonic pain (smallest cluster-level $p = .1925$). Neither direct tonic-condition contrast reached significance. The separate tonic-by-phasic intensity interaction test also did not reach significance (smallest cluster-level $p = .1575$). Correction was performed separately for each contrast and power representation.

### 4 LINEAR MIXED MODEL OUTPUTS

We provide detailed LMM outputs and comparisons of random-effects specifications in this section.

We refitted choice distance bias and SCR associations with identical observations and fixed effects, retaining subject-specific intercepts and slopes in both specifications. The maximal model (M) was `y ~x + (1 + x | subject)`; the no-correlation model (N) was `y ~x + (1 | subject) + (0 + x | subject)`. Singularity was assessed using `lme4::isSingular`.

**Table S2.** Fixed-effect slopes for maximal (M) and no-correlation (N) models. The primary analyses use N throughout the choice-distance and SCR analysis blocks. Confidence intervals are 95% Satterthwaite-based Wald intervals; $p$ values are two-sided and unadjusted.

| Association | Model | $\beta$ | 95% CI | $t$ (df) | $p$ | Singular |
|---|---|---|---|---|---|---|
| *Experiment 1: choice distance bias (Figure 3B)* | | | | | | |
| Distance–rating | M | 0.01072 | $[0.00033, 0.02111]$ | $2.14(22.62)$ | .0438 | Yes |
| | N | 0.01088 | $[0.00134, 0.02042]$ | $2.29(53.05)$ | .0262 | No |
| *Experiment 1: SCR* | | | | | | |
| Fixation–decision value | M | $-0.01143$ | $[-0.02489, 0.00203]$ | $-2.11(5.65)$ | .0823 | No |
| | N | $-0.00802$ | $[-0.01910, 0.00307]$ | $-1.72(6.79)$ | .1302 | No |
| Fixation–rating | M | 0.00661 | $[-0.00572, 0.01893]$ | $1.14(15.28)$ | .2715 | No |
| | N | 0.00649 | $[-0.00566, 0.01864]$ | $1.13(16.20)$ | .2743 | No |
| Shock–decision value | M | $-0.01042$ | $[-0.05333, 0.03250]$ | $-0.65(4.49)$ | .5499 | Yes |
| | N | $-0.01019$ | $[-0.04240, 0.02202]$ | $-0.73(8.08)$ | .4870 | No |
| Shock–rating | M | 0.04338 | $[0.01309, 0.07367]$ | $3.05(14.91)$ | .0081 | Yes |
| | N | 0.03906 | $[0.01470, 0.06343]$ | $3.38(17.02)$ | .0035 | No |
| *Experiment 2: SCR* | | | | | | |
| Fixation–decision value | M | $-0.00072$ | $[-0.00187, 0.00044]$ | $-1.32(15.35)$ | .2057 | No |
| | N | $-0.00073$ | $[-0.00184, 0.00038]$ | $-1.37(18.96)$ | .1857 | No |
| Fixation–rating | M | 0.00100 | $[0.00023, 0.00177]$ | $2.73(18.29)$ | .0137 | No |
| | N | 0.00095 | $[0.00017, 0.00173]$ | $2.54(18.78)$ | .0202 | No |
| Shock–decision value | M | $-0.01224$ | $[-0.02005, -0.00443]$ | $-3.32(16.01)$ | .0043 | No |
| | N | $-0.01364$ | $[-0.02062, -0.00666]$ | $-4.26(11.86)$ | .0011 | No |
| Shock–rating | M | 0.01056 | $[0.00769, 0.01342]$ | $7.62(22.50)$ | $< .001$ | No |
| | N | 0.01071 | $[0.00779, 0.01363]$ | $7.59(22.72)$ | $< .001$ | No |

### 5 POSSIBLE NUMERICAL DIFFERENCES IN LARGE LMM FITTING ACROSS DIFFERENT HARDWARE PLATFORMS

No significant numerical differences were found between AMD and Intel platform for Fig. 10 results. They were reported to differ across platforms in previous version of this paper. The improved numerical stability could be attributed to model reduction: the correlation between random effects was removed from the original maximal model. However, small numerical differences are still possible for mixed model fitting results depending on the hardware platform and environment.

### 6 DISCUSSION OF PAIN INTENSITY INFORMATION AND MODEL ROBUSTNESS

#### 6.1 The Information Lag Caveat

In Experiment 1, the specific pain intensity associated with green pineapples ($x_{stim}$) was held constant within a block but was not explicitly signaled to the participant at the block's onset. Consequently, the precise value of $x_{stim}$ only became known to the agent after the first green pineapple was harvested. In our primary model-fitting, we assumed a constant $x_{stim}$ for the entire one-minute block.

We acknowledge that this is a simplification. Theoretically, the decisions made *before* the first green pineapple pickup might be driven by an expectation of pain rather than the experienced value. However, we opted for the constant $x_{stim}$ approach for several practical and statistical reasons:

- **Data Integrity and Imbalance:** Discarding the pre-exposure period would result in variable analysed durations across subjects and conditions. Among the 20 participants included in the model analysis, 15 of 200 blocks (7.5%) contained exactly one green choice scored by the model. Trimming through that choice would leave only yellow selected outcomes in those blocks, reducing observations of painful choices while retaining avoidance comparisons.

- **Empirical Validation:** We conducted a sensitivity analysis by fitting the model exclusively to data following the first green fruit pickup. The qualitative results and statistical significance of the effort-pain trade-offs remained identical to those reported in the main text (Fig. S8, S9).

In Experiment 2, when no green fruit was picked up in a block, the model retained the most recently experienced shock intensity as a proxy for expected pain, because the current block's intensity had not been sampled; this occurred in one participant's first analysed no-pain block, for which $x_{\mathrm{stim}} = 1/3$. This carry-over rule is a simplifying assumption about unobserved expectations, and in the model with separate vigour constants it also assigns that block the low-pain $C_v$ despite its nominal no-pain condition.

##### 6.2 Future Design Considerations

We explored several alternative designs to resolve this information lag, which may inform future studies:

1. **Pre-block Stimulation:** Providing a sample pulse before the block starts. In pilot testing, we found that stationary stimulation (at rest) was perceived differently from stimulation received during the high-exertion foraging task, potentially introducing a different form of bias.

2. **Extended Block Duration:** Increasing the block length beyond 60 seconds would provide more post-exposure data. However, the physical demands of the VR task introduce fatigue-related performance decay; our pilot data suggested a 30-minute limit for continuous active VR to maintain participant safety and data consistency.

3. **Stochastic RL Frameworks:** Transitioning to a free-operant multi-bandit task with stochastic outcomes would allow for traditional reinforcement learning (RL) updates. However, the increased learning complexity might mask the primary avoidance and effort-based trade-offs we aimed to isolate in this study.

In summary, while the 'information lag' exists, our validation analyses suggest it does not undermine the evidence supporting underlying motivational trade-offs.

#### 7 CLOCK SYNCHRONISATION

##### 7.1 Affine correction of the wireless sensor clock

The wireless skin-conductance recordings in Experiment 1 used an Arduino clock that was independent of the host-computer clock used for task-event timestamps. The difference between these clocks increased during recording, so a constant offset estimated near the beginning of a block did not account for the accumulating timing discrepancy. We therefore estimated both the relative clock rate and the offset separately for each 60-s block.

For packet $i$ received during block $b$, let $u_{bi}$ be the Arduino timestamp of its last sample, in microseconds, and $r_{bi}$ the host timestamp at packet receipt, in milliseconds. We converted the local timestamps to milliseconds, $a_{bi} = u_{bi}/1000$, and fitted the affine relationship by ordinary least squares:

$$(\widehat{\alpha}_b, \widehat{\beta}_b) = \arg\min_{\alpha,\beta} \sum_{i \in \mathcal{P}_b} \left[r_{bi} - (\alpha a_{bi} + \beta)\right]^2 , \tag{1}$$

where $\mathcal{P}_b$ contains packets received between the block start and 60 s later. Each sample's Arduino timestamp $u$ was then placed on the host time axis as

$$\widehat{t}_{\mathrm{host}}(u) = \widehat{\alpha}_b \frac{u}{1000} + \widehat{\beta}_b. \tag{2}$$

The scale factor $\widehat{\alpha}_b$ corrects the relative clock rate, while $\widehat{\beta}_b$ accounts for the offset between the clock origins together with any average unmeasured receipt delay. This mapping was applied before event-aligned extraction and resampling; task-event timestamps remained on the host clock. The same block-specific mapping was used for the SCR averages and the time series fitted to canonical response functions. Fits were rejected when fewer than two packets were available or a backward jump in the local packet-end timestamps occurred within the fitting window. Estimating separate maps for each block avoided fitting across clock resets in inter-block gaps.

Linear clock mapping is also used in offline Lab Streaming Layer (LSL) workflows: the reference XDF importer fits clock-offset measurements to account for linear drift (Kothe et al., 2025). LSL obtains those measurements through repeated two-way network exchanges, whereas our mapping used one-way packet-receipt timestamps. The methods therefore share a regression-based correction principle without implying equivalent timing accuracy. In particular, clock mapping does not independently calibrate delays between physical acquisition and timestamping. Such delays also require separate assessment in LSL systems, for example using a common hardware trigger recorded by both systems (Kothe et al., 2025). Our checks below assess relative alignment and drift reduction, rather than absolute acquisition-to-event latency.

##### 7.2 Evidence for improved temporal alignment

We examined the sensor-to-PC clock relationship, including recordings from participants omitted from the final SCR analyses. In recordings without backward clock jumps, offsets estimated from the first 5 s of each block changed progressively across the recording (Fig. S10A,B). Packet offsets also changed within individual blocks (Fig. S10C). These are two views of the same accumulating timing discrepancy, consistent with a relative clock-rate mismatch. A constant offset aligns the clocks locally; the block-specific affine mapping additionally accounts for their relative rate. Both views are based on recorded timestamps and do not constitute an independent physiological validation. As a descriptive waveform check, we grouped pick-up-aligned SCRs into successive 15-s within-block intervals and cross-correlated each mean post-event waveform with one fixed template: the grand mean of the affine-corrected epochs (Fig. S10D). The same template was used for every interval and both clock mappings, so the comparison concerns changes in relative lag rather than absolute SCR peak latency. The lag trend was flatter after affine correction, consistent with reduced within-block timing drift. However, the template was derived from the corrected data, and overlapping responses and finite-window correlation can affect the estimated lags or attenuate small shifts; flatness therefore does not establish that residual drift is absent.

##### 7.3 EEG trigger pairing and affine clock correction

For the LiveAmp EEG recordings in Experiment 2, we estimated the relationship between the PC clock and the EEG sample clock using triggers recorded in both systems. The acquisition program stored a PC timestamp for each trigger command, and a Wireless Trigger system (Brain Products GmbH) transmitted the trigger to the amplifier, where its marker was recorded with the EEG samples. The manufacturer specifies a typical transmission delay of $2.5 \pm 0.1$ ms, jitter of $\pm 1$ sample point, and a typical trigger loss rate below $1\%$, depending on the environment (Brain Products GmbH, 2020).

The PC sequence comprised the recorded `last_trigger_on_client` values. The EEG sequence comprised the retained hardware markers S2–S61 and S1, including S1 markers renamed by experimental condition during preprocessing that were used to generate ERP figure directly (Fig. 6); their sample positions were preserved. Let $t_1, \ldots, t_N$ denote the ordered PC timestamps in milliseconds, and $n_1, \ldots, n_M$ the ordered EEG trigger positions in samples, calculated from annotation onsets at the nominal sampling frequency $f_s = 500$ Hz.

We considered every feasible initial correspondence $j_1 \in \{1, \ldots, N - M + 1\}$. For each starting point, subsequent EEG triggers were paired greedily to the later PC timestamp closest to $t_{j_{i-1}} + 1000(n_i - n_{i-1})/f_s$, while maintaining increasing PC indices and leaving enough PC entries to complete the pairing. This procedure could skip PC records when an EEG trigger was missing. For each complete candidate index sequence $\mathbf{j} = (j_1, \ldots, j_M)$, we fitted an affine relationship by ordinary least squares and calculated

$$Q(\mathbf{j}) = \min_{a,b} \sum_{i=1}^{M} \left[n_i - (a\, t_{j_i} + b)\right]^2 . \tag{3}$$

The candidate with the smallest summed squared residual was retained. The search thus covered all feasible first correspondences, with one greedy continuation per starting point; it was not an exhaustive enumeration of every possible trigger subset.

The selected coefficients mapped a PC event timestamp $t$ directly to a sample position in the cropped, cleaned EEG:

$$\widehat{n}(t) = \widehat{a}\, t + \widehat{b}. \tag{4}$$

Here $\widehat{a}$ is measured in samples per millisecond and is close to the nominal value 0.5; $\widehat{b}$ includes the recording's sample origin and crop offset. The implementation centred PC timestamps during regression for numerical stability. The same recording-specific map was used to locate PC-timestamped analysis events and block boundaries; event locations were converted to integer sample indices without resampling the EEG.

In the acquisition program, the PC timestamp was captured after writing the trigger value to the serial port and requesting a 10-ms thread sleep, before resetting the trigger value to zero. The fitted offset therefore included the net difference between this software timestamp and hardware marker arrival, including transmission and scheduling delays. The fit corrected relative clock offset and rate, but did not independently calibrate the latency between a physical event and its PC timestamp.

##### 7.4 EEG clock-fit diagnostics

Figure S11 shows the fitted clock relationships and residuals for all 31 available cleaned Experiment 2 recordings before downstream participant exclusions. The upper panel expresses the difference between nominal EEG time and PC time relative to the first matched trigger, $2(n_i - n_1) - (t_{j_i} - t_{j_1})$, in milliseconds. The lower panel shows the residuals $2[n_i - \widehat{n}(t_{j_i})]$ in milliseconds. These residuals reflect timing jitter and any remaining deviation from a constant relative clock rate. The diagnostic supports the selected correspondence and improved relative alignment; it does not uniquely identify the source of clock-rate changes or establish absolute acquisition-to-event latency.

#### 8 SKIN CONDUCTANCE SENSITIVITY ANALYSES

We examined whether the associations reported in the main-text SCR analysis were sensitive to the treatment of uncertainty in the block-level coefficient estimates, and whether they were also supported in Experiment 2. Each experiment was analysed separately. In Experiment 2, two of the 27 participants retained after eye-tracking exclusions were additionally excluded from the SCR analyses because manual inspection identified low-quality recordings, leaving 25 participants. All three approaches used the same outcome-specific observations, raw SCR coefficients, and standardised predictors within each experiment. No first-level significance threshold was applied; the absolute-$z$ exclusion described in Methods was retained.

The primary analysis used unweighted linear mixed-effects models with uncorrelated subject-specific intercepts and slopes throughout the SCR analysis block. The supplementary models below used the same no-correlation specification. Two supplementary inverse-variance mixed-effects approaches incorporated the estimated sampling uncertainty of each block's coefficient. Standard errors were calculated from the first-level ordinary least-squares fits using a Newey–West heteroskedasticity- and autocorrelation-consistent (HAC) covariance estimate, with automatic bandwidth selection, first-order prewhitening, and a finite-sample adjustment (Newey and West, 1987, 1994). The frequentist analysis used `metafor` to fit a multilevel Gaussian model by restricted maximum likelihood (Viechtbauer, 2010). The supplied sampling variances were the squared HAC standard errors. Subject-specific intercepts and slopes accounted for participant variation, and an additional observation-level random intercept accounted for residual block heterogeneity. Fixed-effect inference used participant-clustered CR2 standard errors and Satterthwaite degrees of freedom through `clubSandwich` (Pustejovsky and Tipton, 2018).

The Bayesian analysis used `brms` with the same Gaussian observation model and subject-level structure (Bürkner, 2017). The formula's `se(..., sigma=TRUE)` term combines the supplied squared HAC standard error with an estimated residual variance. Both approaches account for coefficient-estimation uncertainty conditional on the supplied HAC sampling variances, which were treated as fixed inputs; uncertainty in those variance estimates themselves was not propagated.

Broad proper priors were retained: a normal prior with mean zero and standard deviation 1 for the standardised-predictor slope, and positive-truncated Student-$t$ priors with 3 degrees of freedom, location zero, and scale 2.5 for subject-level standard deviations and the residual standard deviation. The default intercept prior was Student-$t$ with 3 degrees of freedom and scale 2.5; its location was 0.1 for the Experiment 1 shock models and zero for the remaining models. Four chains were run for 4,000 iterations each, including 1,000 warmup iterations. Bayesian point estimates are posterior medians, with equal-tailed 95% credible intervals; sign probabilities are Monte Carlo estimates, and rounding to 1 does not imply certainty. Frequentist intervals are 95% confidence intervals and $p$ values are two-sided and unadjusted.

The positive shock–rating association was supported by all three approaches in both experiments (Tables S3 and S4). The fixation–decision-value association was not established in either experiment. Fixation–rating associations were sensitive to the analysis approach: the Experiment 1 METAFOR result and the Experiment 2 unweighted result met the nominal significance threshold, whereas the other confidence or credible intervals included zero. Shock-evoked coefficients were negatively associated with decision values across all three approaches in Experiment 2, but this association was not established in Experiment 1. With the model's coding, more negative decision values represent more aversive pain. The shock–rating association was therefore the only association supported across both experiments by all three approaches. This pattern does not establish either a difference between the associations or an absence of the associations whose intervals included zero.

**Table S3.** SCR association estimates in Experiment 1. Slopes are in raw SCR-coefficient units per predictor standard deviation. Bracketed intervals are 95% confidence intervals for the frequentist models and equal-tailed credible intervals for BRMS. The Bayesian probability is the posterior probability of the displayed effect direction.

| Association | Unweighted LMM | METAFOR + CR2 | BRMS |
|---|---|---|---|
| Fixation–decision value | $-0.00802$<br>$[-0.01910, 0.00307]$<br>$p = 0.1302$ | $-0.00638$<br>$[-0.01432, 0.00157]$<br>$p = 0.1005$ | $-0.00716$<br>$[-0.01691, 0.00050]$<br>$P(\beta < 0) = 0.9662$ |
| Fixation–rating | 0.00649<br>$[-0.00566, 0.01864]$<br>$p = 0.2743$ | 0.00600<br>$[0.00002, 0.01199]$<br>$p = 0.0495$ | 0.00570<br>$[-0.00188, 0.01211]$<br>$P(\beta > 0) = 0.9349$ |
| Shock–decision value | $-0.01019$<br>$[-0.04240, 0.02202]$<br>$p = 0.4870$ | $-0.02089$<br>$[-0.05021, 0.00844]$<br>$p = 0.1446$ | $-0.02697$<br>$[-0.08916, 0.00565]$<br>$P(\beta < 0) = 0.9483$ |
| Shock–rating | 0.03906<br>$[0.01470, 0.06343]$<br>$p = 0.0035$ | 0.04914<br>$[0.02424, 0.07403]$<br>$p < .001$ | 0.04879<br>$[0.02450, 0.07242]$<br>$P(\beta > 0) = 0.9998$ |

**Table S4.** SCR association estimates in Experiment 2. Slopes are in raw SCR-coefficient units per predictor standard deviation. Bracketed intervals are 95% confidence intervals for the frequentist models and equal-tailed credible intervals for BRMS. The Bayesian probability is the posterior probability of the displayed effect direction.

| Association | Unweighted LMM | METAFOR + CR2 | BRMS |
|---|---|---|---|
| Fixation–decision value | $-0.00073$<br>$[-0.00184, 0.00038]$<br>$p = 0.1857$ | $-0.00027$<br>$[-0.00085, 0.00030]$<br>$p = 0.3128$ | $-0.00029$<br>$[-0.00109, 0.00048]$<br>$P(\beta < 0) = 0.7853$ |
| Fixation–rating | 0.00095<br>$[0.00017, 0.00173]$<br>$p = 0.0202$ | 0.00036<br>$[-0.00005, 0.00078]$<br>$p = 0.0826$ | 0.00038<br>$[-0.00006, 0.00087]$<br>$P(\beta > 0) = 0.9570$ |
| Shock–decision value | $-0.01364$<br>$[-0.02062, -0.00666]$<br>$p = 0.0011$ | $-0.00945$<br>$[-0.01376, -0.00513]$<br>$p < .001$ | $-0.00958$<br>$[-0.01490, -0.00540]$<br>$P(\beta < 0) = 0.9999$ |
| Shock–rating | 0.01071<br>$[0.00779, 0.01363]$<br>$p < .001$ | 0.00889<br>$[0.00606, 0.01173]$<br>$p < .001$ | 0.00889<br>$[0.00619, 0.01187]$<br>$P(\beta > 0) = 1.0000$ |

### 9 GRID SEARCH PARAMETER CONFIGURATION

Grid search for hyperparameters that best fit the model to the experimental data is deterministic. The key information to reproduce the fitting results is the fitting grid. We present the grid information below.

Experiment 1 (Figure 3)

A two level hierarchical grid search to reduce the search space while maintaining high resolution output. The algorithm is carried out by library functions provided by Scipy package, specifically, the scipy.optimize.brute function.

First level:

```
x scale in u: slice(1, 10.1, 1),
x translation in u: slice(-10, 10.1, 0.5),
positivity of C_p: slice(-1, 1.1, 2),
C_m horizontal: slice(-1, 1.01, 0.1),
C_m vertical: slice(-1, 1.01, 0.1)
```

`slice` is a builtin python function which specifies starting point, end point (excluded), and steps in a 3-tuple. We use it here to represent the grid parameters. $C_p$ is calculated by $(1 - \|C_m\|_2)$ multiplied by positivity of $C_p$. For grid points with $\|C_m\|_2 > 1$, the objective function returned the maximum error rate of 1 to exclude these cases.

Let `res` be the tuple returned from first level, we have the following grid for the second level:

```
slice(res[0] - 0.9, res[0] + 0.91, 0.1),
slice(res[1] - 0.5, res[1] + 0.51, 0.01),
{res[2]},
slice(res[3] - 0.1, res[3] + 0.11, 0.01),
slice(res[4] - 0.1, res[4] + 0.11, 0.01)
```

Experiment 2 (Figure 9)

Single vigour constants ($C_v$) across phasic pain conditions (Figure 9A):

```
x scale in u: {0.125, 0.25, 0.5, 1, 2, 4, 8, 16},
x translation in u: slice(-1, 1, 0.1),
C_v: slice(1, 100, 1),
C_p: slice(-100, 100, 1),
C_m horizontal: slice(0, 1, 0.001)
```

$C_m$ vertical is calculated to satisfy $\|C_m\|_2 = 1$. `slice` here is end-inclusive.

Separate vigour constants ($C_v$) across phasic pain conditions (Figure 9B)

First level:

```
x scale in u: {0.125, 0.25, 0.5, 1, 2, 4, 8, 16},
x translation in u: slice(-1, 1, 0.2),
C_v (no pain): slice(1, 101, 10),
C_v (low pain): slice(1, 101, 10),
C_v (high pain): slice(1, 101, 10),
C_p: slice(-100, 100, 1),
C_m: slice(0, 1, 0.01)
```

Second level:

Let `fitting_result` be the tuple returned from first level, we have the following grid for the second level:

```
{fitting_result[0]},
slice(fitting_result[1] - 0.1f, fitting_result[1] + 0.1f, 0.1),
slice(max(fitting_result[2] - 10, 1.0f), min(fitting_result[2] + 10, 101.0f), 1),
slice(max(fitting_result[3] - 10, 1.0f), min(fitting_result[3] + 10, 101.0f), 1),
slice(max(fitting_result[4] - 10, 1.0f), min(fitting_result[4] + 10, 101.0f), 1),
slice(max(fitting_result[5] - 1, -100.0f),
      min(fitting_result[5] + 1, 100.0f), 0.01),
slice(max(fitting_result[6] - 0.01f, 0.0f),
      min(fitting_result[6] + 0.01f, 1.0f), 0.0001)
```

### 10 EYE TRACKING METHODS AND GENERATION ATTEMPT

#### 10.1 Acquisition and detection of stalled data

Eye-tracking data were acquired from the HTC Vive Pro Eye through the Vive SRanipal SDK using an asynchronous callback. Unity accessed the most recently received eye-data record on each rendering frame and used the gaze direction to cast a ray from the virtual camera position into the scene. The object intersected by this ray was recorded as `gaze_object`; this object sequence supplied the gaze-dependent observation and decision events used in the analyses. Throughout, fixation on an object refers to intersection of the gaze ray with that object.

The acquisition code checked the validity of the stored eye data but did not check whether its frame number had advanced. When the callback stopped supplying new data, the previous record could therefore remain marked as valid, and Unity continued raycasting with the stale `gaze_direction`. Head movements and changes in the scene could still change the raycast object despite the unchanged eye-data record. Initial analyses used the object sequence rather than inspecting the underlying gaze vectors, which obscured this acquisition failure.

We subsequently screened the analysed 60-s blocks. Within each block, the script counted exact occurrences of each recorded gaze-direction vector and flagged a participant if any vector occurred in more than 100 frames. Four participants in each experiment met the criterion and were excluded from analyses requiring eye tracking, leaving 20 participants in Experiment 1 and 27 in Experiment 2. Exclusions applied at the participant level, including participants whose failure affected only some analysed blocks. Participants whose eye-tracking failure occurred only outside the analysed blocks were retained in the study analyses but excluded from gaze-generation model training. Gaze-dependent analyses included choice-distance trade-off estimates, computational-model fits and their downstream analyses, and physiological responses aligned to gaze-derived events.

The acquisition code attempted to obtain combined gaze before falling back to the left or right eye, but the original recordings did not retain which source supplied each gaze ray. Later diagnostic captures, which additionally recorded these separate outputs, were consistent with the recorded direction being a combined gaze vector formed from a convex mixture of the left- and right-eye directions. The empirical relationship between mixture weights and eye openness suggested confidence-related weighting, but the exact weighting algorithm was unknown. These observations do not establish the source selected on each original frame or independently verify the SDK's combination rule.

#### 10.2 Exploratory generation of gaze from head movements

Because gaze-dependent observations were required for several analyses, we explored offline generation from head rotation using HAGI (Jiao et al., 2025b), specifically the rotation-only HAGI++ imputation model (Jiao et al., 2025a). A pretrained checkpoint was fine-tuned on recordings from 18 participants in Experiment 1, using no Experiment 2 recordings for fine-tuning. The model was a transformer-based diffusion model with four layers, eight attention heads, a hidden dimension of 64, and 50 diffusion steps. Fine-tuning used AdamW for 1,000 epochs, with learning rate $2 \times 10^{-4}$, batch size 256, weight decay $10^{-6}$, and random seed 0; there was no separate validation split. These settings describe the exploratory model rather than a model selected through held-out validation.

Head rotations were resampled to 30 Hz using quaternion interpolation and represented by relative rotation matrices between consecutive samples. Training used five-second windows (150 samples) with 50% overlap and at least 95% observed gaze, drawn from the full available recordings rather than only the analysed task blocks. Gaze targets were represented by the sine of head-relative pitch and yaw. At generation, all gaze samples were masked, so the model received head rotation but no observed gaze, wrist motion, or visual scene information. The imputation-trained model was thus applied to the full-generation case. Inference used non-overlapping five-second windows, with the componentwise median of 20 diffusion samples in angular coordinates converted back to unit directions and interpolated to the original frame timestamps. This was an offline procedure with access to the head-motion sequence within each window.

We evaluated directional agreement on the 27 Experiment 2 participants who passed the original-data quality screen, using their 18 analysed blocks each. Recorded and generated directions were compared in the head-relative coordinate system, independently of Unity raycasts. Frame timestamps had to match exactly, and only nonzero vectors from both sources were eligible. We normalised both vectors and calculated their angular separation as $\arccos(\hat{\mathbf{g}}_{\text{recorded}} \cdot \hat{\mathbf{g}}_{\text{generated}})$ in degrees. Mean angular errors were averaged within blocks and then within participants; root-mean-square (RMS) errors were calculated from the corresponding block-averaged squared errors. Group summaries gave participants equal weight.

The fixed baseline was the normalised mean recorded direction from the analysed task blocks of the same 18 Experiment 1 training participants, with blocks and participants weighted equally. It therefore used neither Experiment 2 gaze nor head motion. We additionally calculated each evaluation participant's own mean direction over their analysed blocks as an optimistic descriptive reference; this reference used the evaluation data and was not an independently trained predictor. Generated gaze reduced mean angular error from $11.32 \pm 3.25°$ for the fixed baseline to $9.15 \pm 2.75°$ (mean ± between-participant SD), a 19.2% reduction in the group mean. The participant-level paired reduction was 2.17° (95% CI $[1.85, 2.49]$, two-sided paired $t(26) = 13.94$, $p < .001$). The participant-mean reference had an error of $9.68 \pm 1.98°$; improvement relative to this reference was smaller and uncertain (0.54°, 95% CI $[-0.08, 1.15]$, $t(26) = 1.80$, $p = .084$). Table S5 reports both error measures. These exploratory intervals and tests used participants as the independent units and were not adjusted for multiple comparisons.

**Table S5.** Angular deviation from recorded gaze in Experiment 2 ($n = 27$). Entries are the mean ± between-participant SD of participant-level errors in degrees. Blocks and participants receive equal weight. The training-mean baseline uses only Experiment 1 data; the participant mean uses evaluation data and is a descriptive reference rather than an independent predictor.

| Direction estimate | Mean angular error | RMS angular error |
|---|---|---|
| HAGI++, head rotation only | $9.15 \pm 2.75$ | $10.51 \pm 2.84$ |
| Experiment 1 mean direction | $11.32 \pm 3.25$ | $12.94 \pm 3.34$ |
| Participant mean (evaluation data) | $9.68 \pm 1.98$ | $11.33 \pm 2.14$ |

To compare temporal variation, we converted each unit direction to azimuth $\operatorname{atan2}(x, z)$ and elevation $\operatorname{atan2}(y, \sqrt{x^2 + z^2})$ in degrees. Azimuth was unwrapped within each contiguous segment. Both signals used identical segments and time grids: invalid vectors, duplicate timestamps, and gaps exceeding 100 ms broke segments. We linearly interpolated to 120 Hz and applied the same zero-phase polyphase finite-impulse-response anti-alias filter (Kaiser window, $\beta = 5$) before downsampling to 30 Hz, matching the generator's sampling rate. We discarded 0.5 s from each segment end and estimated power spectral density (PSD) using 6.4-s Hann windows with 50% overlap and constant detrending. Azimuth and elevation PSDs were summed; segment estimates were weighted by their number of Welch windows, then averaged equally across blocks and participants. All 486 blocks contributed spectral estimates. We compared 5–15 Hz power within the shared sampling bandwidth; this descriptive band was not a saccade classifier.

Generated trajectories contained substantially less rapid variation (Fig. S12): mean 5–15 Hz angular power was $0.35 \pm 0.11$ versus $4.69 \pm 2.35$ degree$^2$ for recorded gaze. The mean within-participant generated-to-recorded power ratio was $8.23 \pm 2.31\%$. High-frequency power accounted for $0.84 \pm 0.29\%$ of generated power and $4.30 \pm 1.45\%$ of recorded power over 0.15625–15 Hz. Reduced high-frequency content is consistent with failure to recover some rapid, potentially salience-driven saccades, but PSD alone cannot distinguish saccades from tracking noise or other fluctuations. The generation pipeline's resampling, median aggregation, and windowing may also contribute to smoothing. These results motivate testing visual scene information as an additional input for this task; they do not establish the physiological origin of the missing variation.

##### 10.3 Replay limitations and interpretation

To evaluate whether generated gaze could recover the object sequences needed for behavioural modelling, we attempted to reconstruct the recorded experiment in Unity and repeat raycasting in the replayed scene. As a control, we also supplied the original recorded gaze directions to the replay; discrepancies in this control cannot be attributed to gaze generation. Replay reproduced gaze-object identification for stationary objects, but discrepancies emerged during fruit pickup, dropping, and throwing (Fig. S13). These interactions depend on the timing of external input and on the physics simulation. The original logger sampled during rendering updates

(`Update`), rather than recording each physics update (`FixedUpdate`); replaying those samples therefore did not reliably reproduce the original timing of input, attachment animations, and subsequent object trajectories.

Because a raycast depends on both gaze and the scene state, these replay discrepancies prevented a clean test of whether generated gaze could recover the object sequences and decision events needed to reproduce the computational-model results and the study's hypothesis tests. Generated gaze was therefore not used to reinstate excluded participants in the reported analyses. Directional improvement over a fixed mean baseline does not establish that the missing task-relevant observations have been recovered.

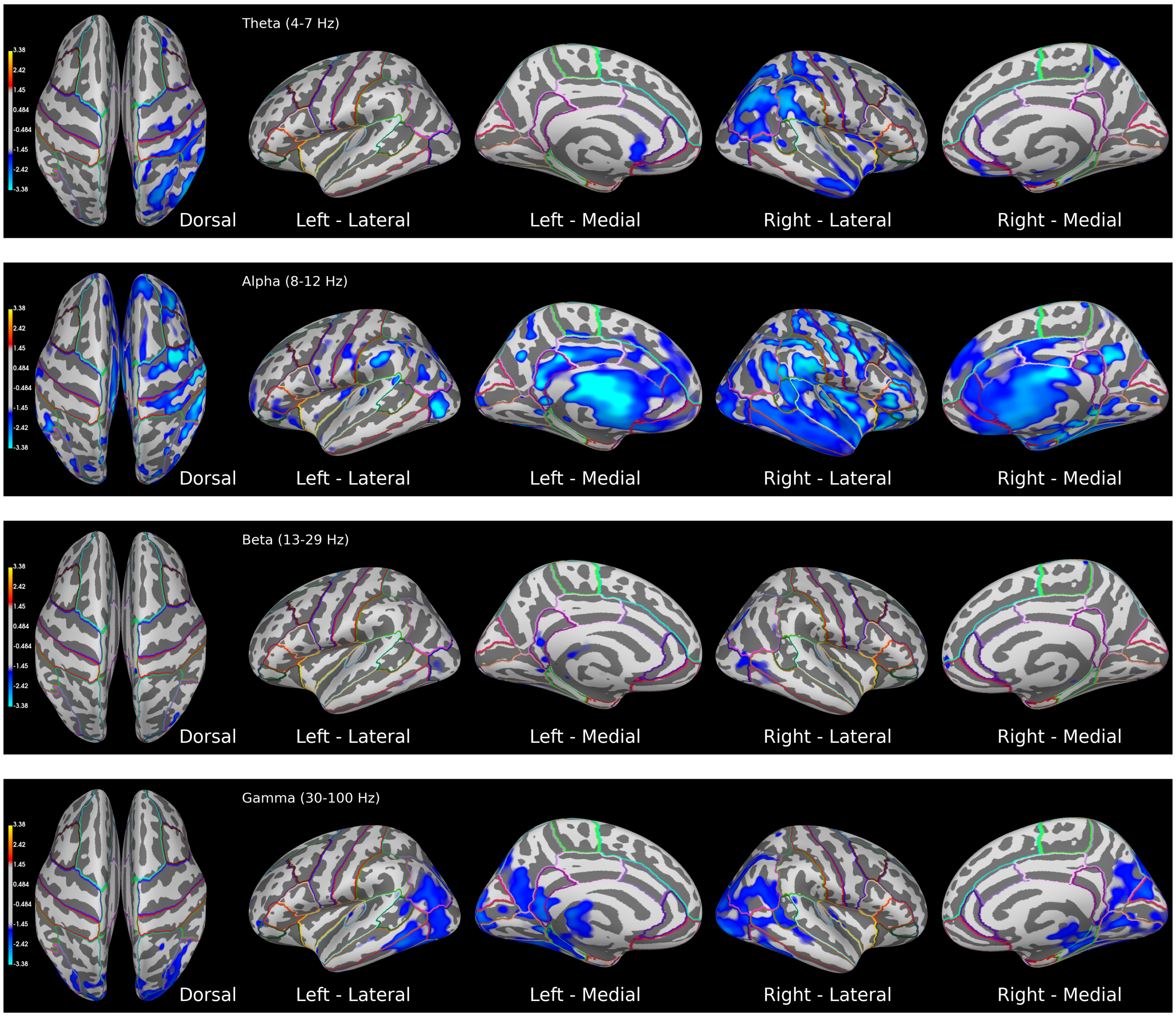

**Figure S1.** t-values for the effect of tonic pain in source space, generated from the LMM. Colours are thresholded at 1.310 for visibility with transparency, 1.697 for non-transparency, and 3.385 for the highest value on the color map. APARC parcellation (Desikan et al., 2006) was overlapped for visual inspection. Frequency band notations represented the pass band used for filtering the signal in each epoch before applying sLORETA in time-domain.

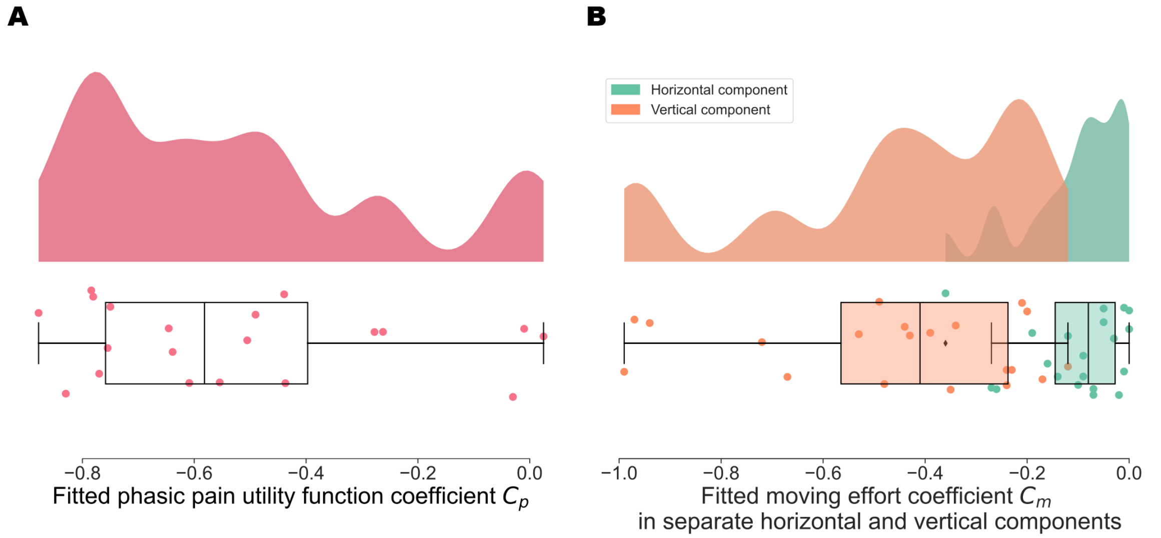

**Figure S8.** Same qualitative result as shown in Fig. 3(C,D). **(A)** The negative phasic pain coefficients ($M = -0.521, SD = 0.275$) showed the model captured the aversiveness of phasic pain stimuli in this free-operant decision-making task, $t(19) = -8.27$, $p < .001$. **(B)** The moving effort coefficient $C_m$ was separated into a horizontal component ($M = -0.105$, $SD = 0.097$) and a vertical component ($M = -0.457$, $SD = 0.265$). The fitted coefficients showed lower effort cost to move horizontally than vertically, $t(19) = 6.03$, $p < .001$.

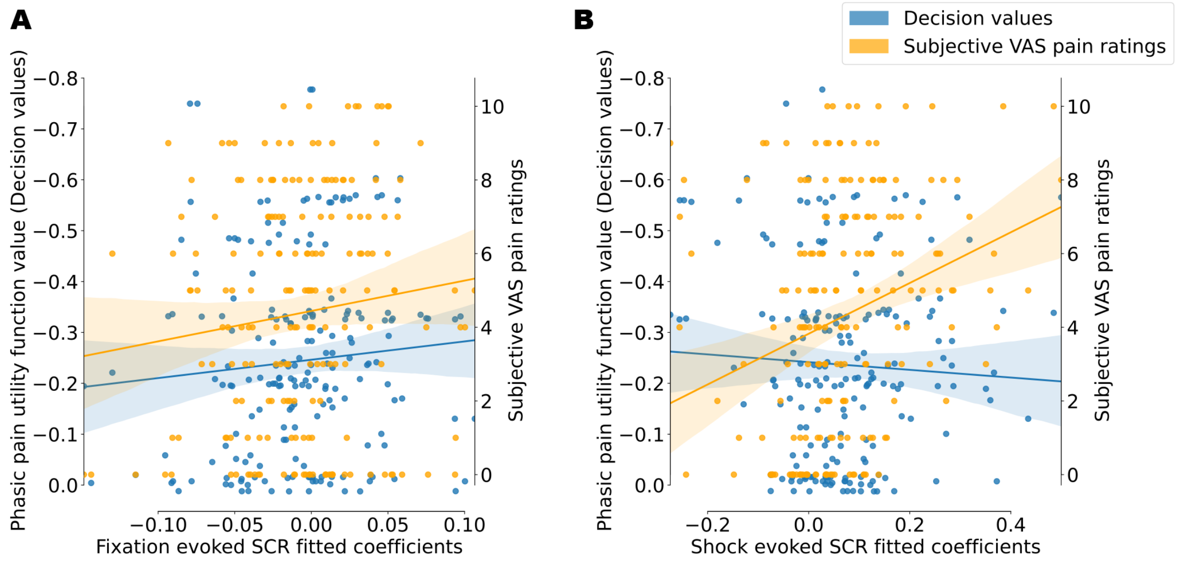

**Figure S9.** Same qualitative result as shown in Fig. 4(B,C). (**A**) In the primary unweighted mixed-effects analyses, the associations of fixation-evoked SCR coefficients with decision values ($\beta = -0.007$, 95% CI $[-0.017, 0.003]$, $t(11.06) = -1.46$, $p = .172$) and subjective ratings ($\beta = 0.0065$, 95% CI $[-0.0057, 0.0186]$, $t(16.20) = 1.13$, $p = .274$) were inconclusive. (**B**) Shock-evoked SCR coefficients were positively associated with subjective ratings ($\beta = 0.0391$, 95% CI $[0.0147, 0.0634]$, $t(17.02) = 3.38$, $p = .0035$); the association with decision values was inconclusive ($\beta = -0.002$, 95% CI $[-0.042, 0.038]$, $t(9.96) = -0.12$, $p = .906$).

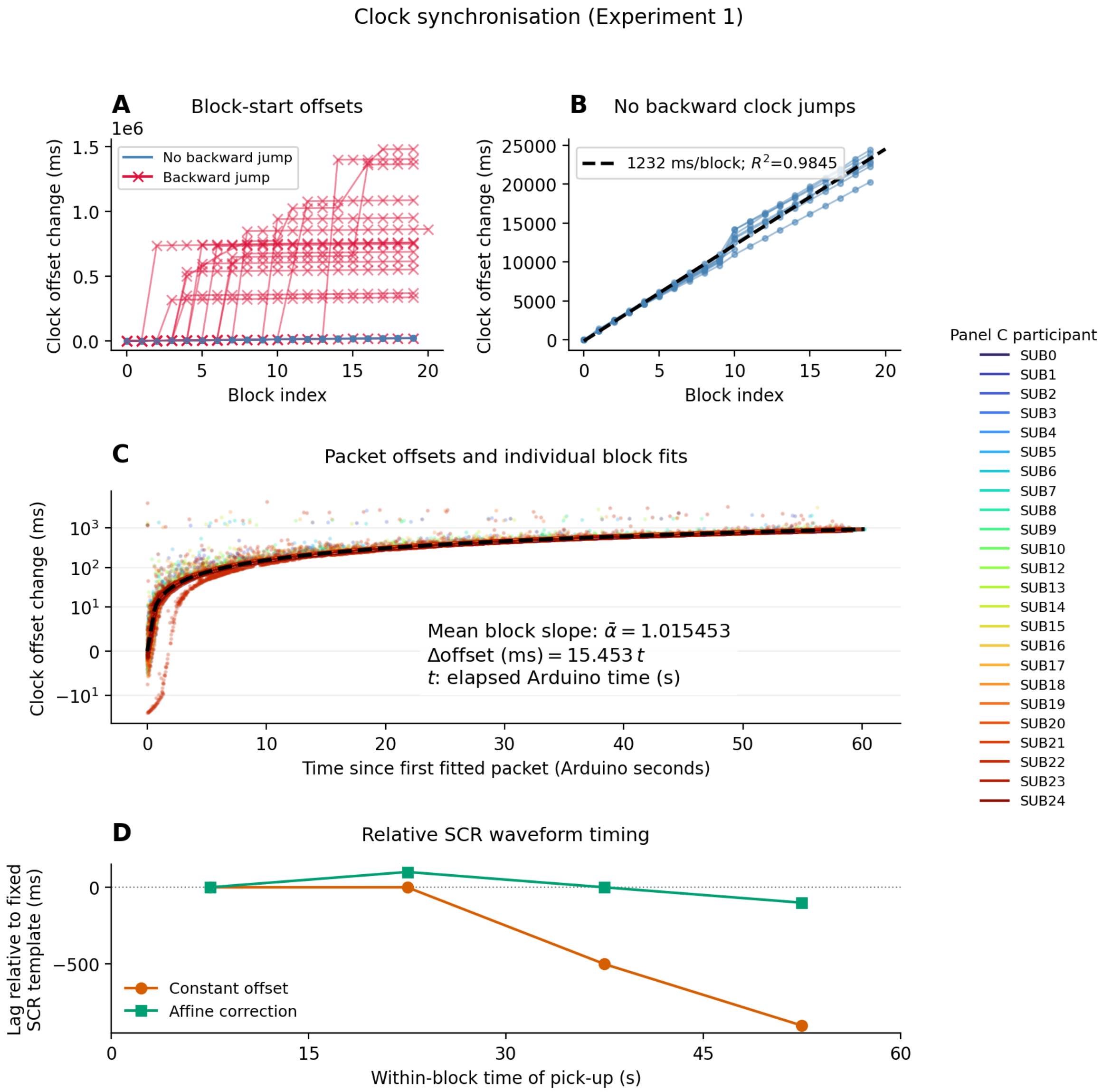

**Figure S10.** Clock synchronisation diagnostics, including participants omitted from the final SCR analyses. **(A)** Block-start offsets relative to the first available offset; red crosses identify recordings with backward clock jumps. **(B)** Recordings without backward jumps, with a descriptive linear fit. **(C)** Packet offsets and block-specific fits, centred at each block's first fitted packet; colours identify participants. The black dashed reference uses the mean block slope, $\bar{\alpha}$: $\Delta o = 1000(\bar{\alpha} - 1)t$, with offset in milliseconds and Arduino time in seconds. Its curvature reflects the signed logarithmic axis. Panels A–C show the same clock-rate discrepancy at different timescales. **(D)** Lags of pick-up-aligned mean SCR waveforms relative to a single fixed template, under each clock mapping. Events include pain-associated fruit pick-ups in zero-stimulation blocks. A flatter trend indicates greater consistency relative to this template, not absolute response latency or proof that small residual drift is absent.

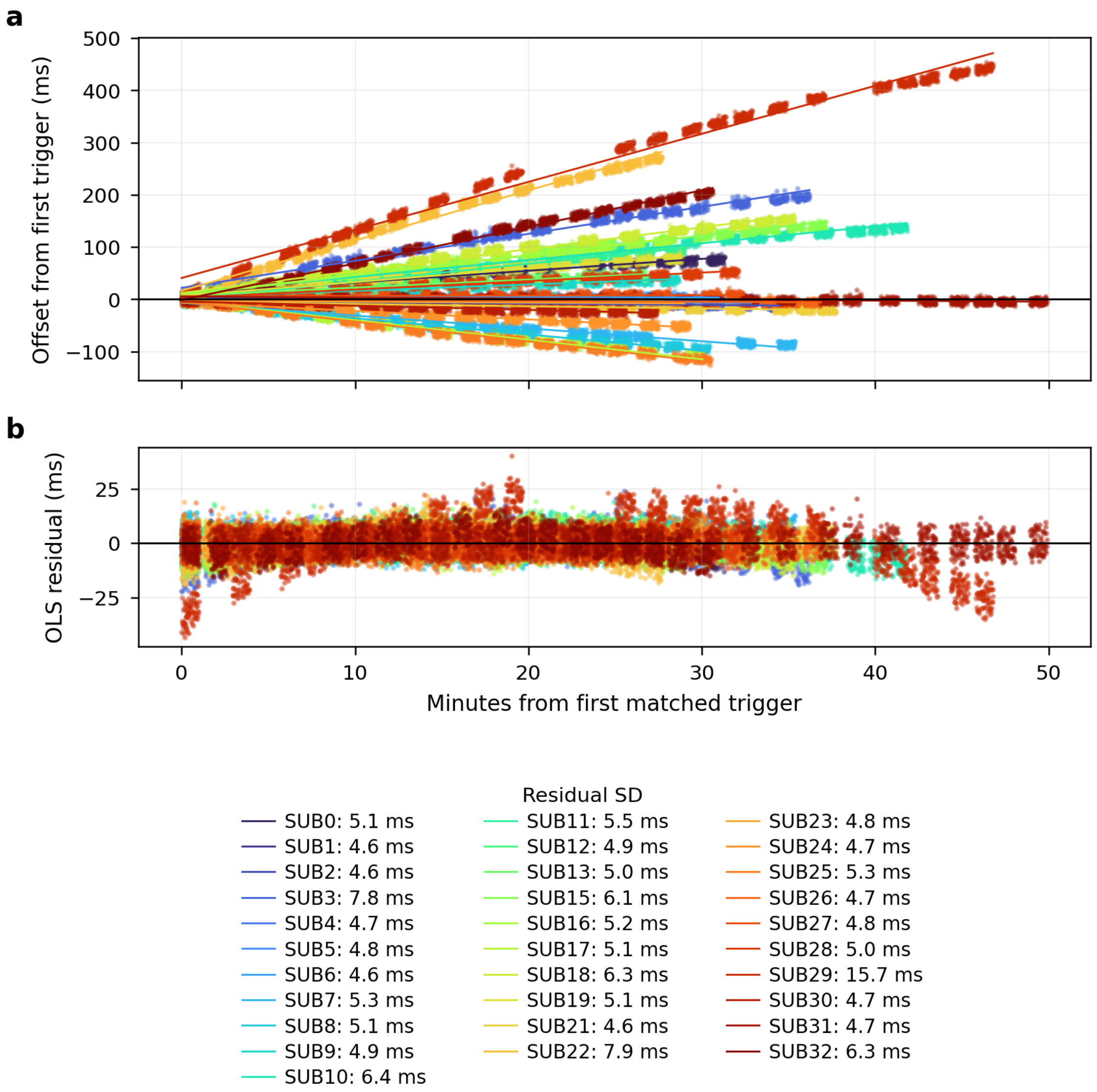

**Figure S11.** EEG–PC clock alignment and residual timing variation in Experiment 2. Colours identify the 31 recordings. **(a)** Differences between nominal EEG time and PC time, relative to the first paired trigger, with the fitted affine relationships superimposed. The changing offsets show the relative clock-rate differences that would remain under a constant-offset correction. **(b)** Residuals of the selected trigger pairs after affine correction, expressed in milliseconds at 500 Hz. The legend reports each recording's residual standard deviation. Most traces have a narrow residual spread; systematic curvature in some recordings, particularly SUB29, indicates remaining departures from a constant relative rate, presumably due to multiple clock skew adjustment on the PC initiated during the data collection period. Residuals include software-timestamp and trigger-delivery variability and should not be interpreted as isolated wireless-transmission jitter.

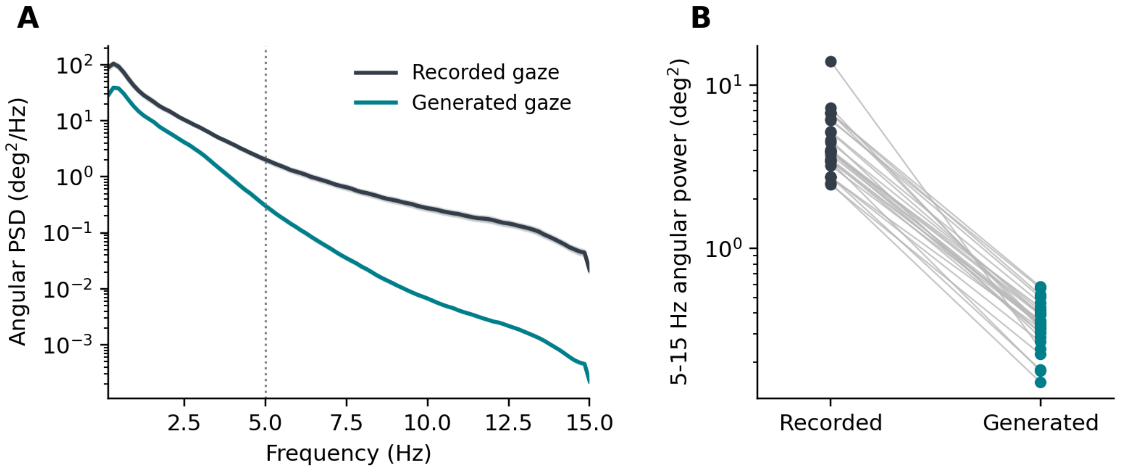

**Figure S12.** Temporal variation in recorded and head-rotation-generated gaze in Experiment 2 ($n = 27$, 18 blocks per participant). **(A)** Sum of azimuth and elevation PSDs, averaged equally across participants after the common preprocessing described in the text. The PSD axis is logarithmic, the frequency axis is linear, and the dotted line marks 5 Hz. **(B)** Integrated 5–15 Hz angular power for each participant; connecting lines identify paired recorded and generated estimates. The lower generated power indicates reduced rapid variation within the shared bandwidth. Constant mean-direction baselines have no temporal power after detrending and are assessed separately in Table S5.

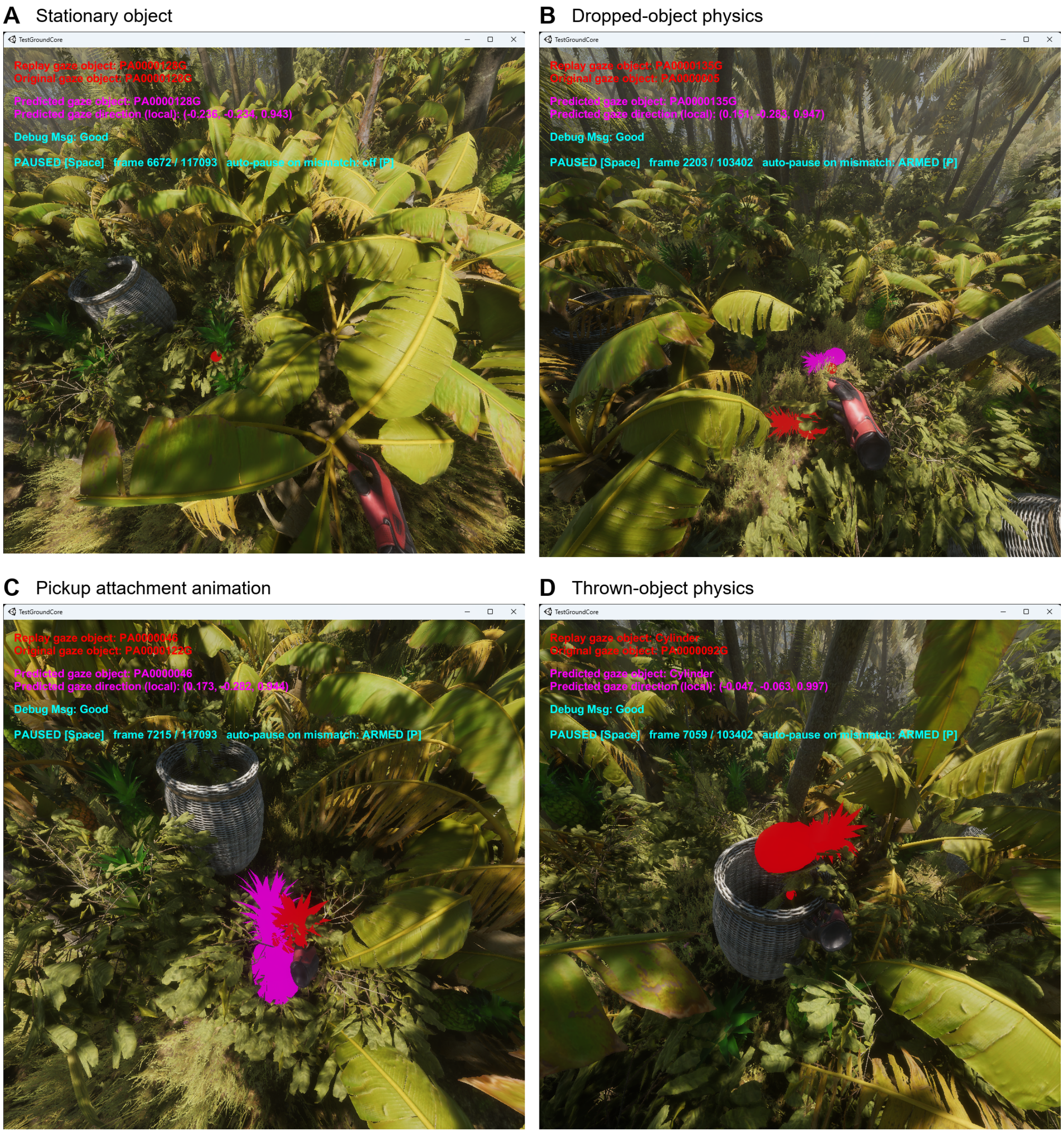

**Figure S13.** Replay validation examples using recorded gaze as the input, illustrating scene-reconstruction limitations independently of generation error. **(A)** A stationary object yielded agreement between the original and replayed gaze-object identifiers. Disagreements occurred during **(B)** dropping, **(C)** pickup attachment animation, and **(D)** throwing. The diagnostic overlays show the recorded object identifier and the object identified by replay raycasting. The screenshots' "Predicted" label denotes the direction supplied to the replay, which in this control reproduces the recorded direction. Raycast disagreement in these moving-object examples prevents attributing differences in downstream results solely to generated gaze.
